# Full site-specific addressability in DNA origami-templated silica nanostructures

**DOI:** 10.1101/2022.12.22.521222

**Authors:** Lea M. Wassermann, Michael Scheckenbach, Anna V. Baptist, Viktorija Glembockyte, Amelie Heuer-Jungemann

## Abstract

DNA nanotechnology allows for the fabrication of nano-meter-sized objects with high precision and selective addressability as a result of the programmable hybridization of complementary DNA strands. Such structures can template the formation of other materials, including metals and complex silica nanostructures, where the silica shell simultaneously acts to protect the DNA from external detrimental factors. However, the formation of silica nanostructures with site-specific addressability has thus far not been explored. Here we show that silica nanostructures templated by DNA origami remain addressable for post silicification modification with guest molecules even if the silica shell measures several nm in thickness. We used the conjugation of fluorescently labelled oligonucleotides to different silicified DNA origami structures carrying a complementary ssDNA handle as well as DNA PAINT super-resolution imaging to show that ssDNA handles remain unsilicified and thus ensure retained addressability. We also demonstrate that not only handles, but also ssDNA scaffold segments within a DNA origami nanostructure remain accessible, allowing for the formation of dynamic silica nanostructures. Finally we demonstrate the power of this approach by forming 3D DNA origami crystals from silicified monomers. Our results thus present a fully site-specifically addressable silica nanostructure with complete control over size and shape.

## 1. Introduction

DNA nanotechnology allows for the bottom-up synthesis of nanometer-sized objects with high precision and selective addressability for guest molecule placement due to the programmable hybridization of complementary DNA strands. This site-specific addressability renders DNA origami nanostructures as “breadboards” for the sub-nm precise placement of various different guest molecules such as nanoparticles (NPs), fluorophores, and proteins.^[1]^ The rational design of DNA-based nanostructures and their modification with different functional molecules enables a great variety of applications ranging from catalysis^[2]^, to biomedicine^[3]^ and materials science^[4]^. DNA nanostructures have also proven to be excellent templates for the formation of complex materials including polymers^[5]^, metals^[6]^ and biominerals like calcium phosphate^[7]^ and silica.^[4, 8]^ This templating approach allows to create inorganic nanostructures with shapes otherwise not obtainable through standard wet-chemical methods.^[4]^ At the same time, coating of DNA nanostructures with calcium phosphate or silica confers a significantly increased degree of stability as the DNA is essentially fossilized.^[7b, 8a–d]^ Different methods of silicification have been reported. In most cases, structures are initially reacted with the cationic pre-cursor N-trimethoxysilylpropyl-N,N,N-trimethylammonium chloride (TMAPS), which electrostatically associates with the phosphate backbone on the DNA nanostructure. The use of (3-Aminopropyl)triethoxysilane (APTES) instead of TMAPS has also been reported.^[8d]^ Silica formation is then initiated through the addition of tetraethyl orthosilicate (TEOS) in a Stöber-like process.^[9]^ Alternatively pre-clusters of TMAPS and TEOS can be formed which subsequently accumulate and polymerize on the phosphate backbone. Such silicified structures were shown to withstand extreme temperatures of up to 1000°C^[8f]^, high pressures^[8a, 8f]^ and degradation by nucleases^[8b]^. We recently also used small angle X-ray scattering (SAXS) to show that even minimal (sub-nm) silica deposition already results in DNA nanostructures that remain stable during prolonged heating.^[8g]^

Generally, it is assumed that during biomineralization processes, DNA is hermetically sealed giving rise to optimal protection. At the same time, shape and size of the DNA nanostructure templates are being retained. However, the highly attractive possibility of also retaining the site-specific addressability of DNA nanostructures for precise guest molecule placement after biomineralization has thus far seldom been explored. In this work we show that, surprisingly, after silicification, single stranded (ss) handles of DNA or peptide nucleic acids (PNA) protruding from a DNA nanostructure remain unsilicified and hence remain accessible for hybridization and further functionalization. Using simple hybridization experiments with fluorescently labelled oligonucleotides (**Scheme 1a**) or DNA-coated gold nanoparticles (Au NPs) (**Figure S6**) as well as DNA PAINT super-resolution microscopy (**Scheme 1b**), we show that structures silicified in solution with minimal silica deposition as well as structures silicified on a surface with a several nm thick shell remain fully site-specifically addressable. Furthermore, by dynamically changing the shape of a silicified 18 helix bundle (18HB) we demonstrate that not only handles protruding from a structure remain accessible, but also ssDNA segments of scaffold within a nanostructure. Finally, we demonstrate the power of the approach for materials science applications by forming open channel 3D DNA origami-silica hybrid crystals using silicified octahedral monomers. Our work thus demonstrates that silica nanostructures templated by DNA origami combine the properties and mechanical resilience of an inorganic material with the programmability and addressability of DNA origami into a new type of fully site-specifically addressable inorganic nanostructure with complete control over size and shape.

**Scheme 1.**
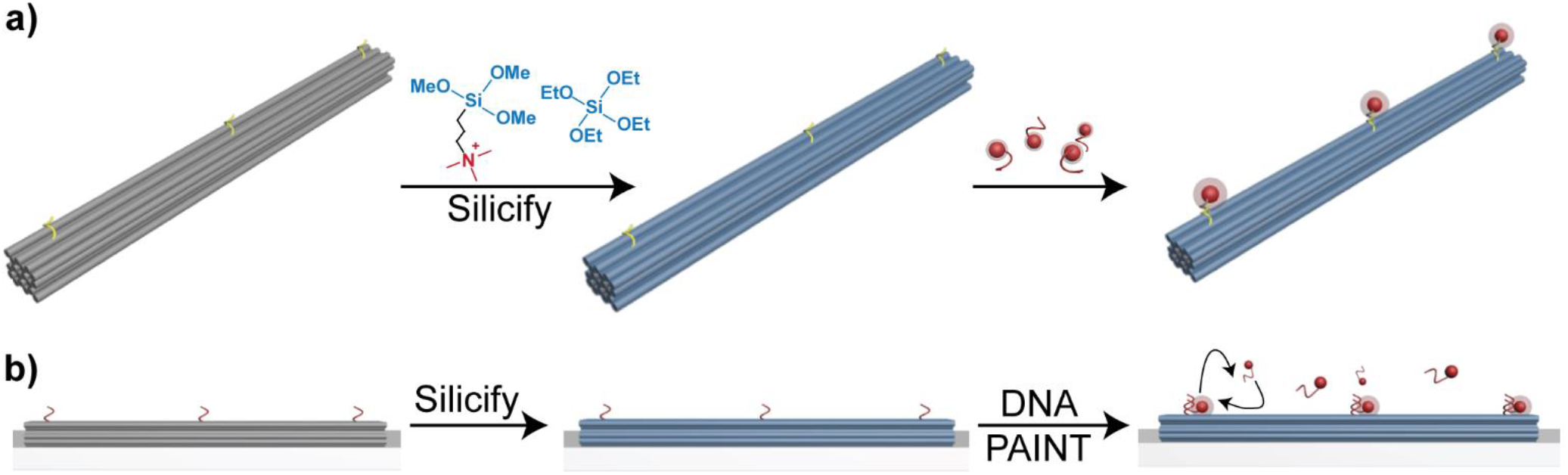
Schematic illustration displaying the assessment of ssDNA handle accessibility on DNA origami after silicification in solution (a) and on a surface (b). Samples silicified in solution are in or near the maximally condensed state and contain a set of ssDNA handles. If these remain unsilicified, a fluorophore-labelled anti-handle will be able to hybridize to the structure. (b) Samples silicified on a surface with silica shell thicknesses of a few nm contain a set of 8 nucleotide (nt) long docking sites to which fluorescently-labelled imager strands bind transiently (DNA PAINT super-resolution microscopy).

## 2. Results and Discussion

As the interaction between DNA and TMAPS/TEOS is based on electrostatic interactions between the anionic phosphate backbone and cationic TMAPS, we initially hypothesized that PNA, with a net neutral charge due to its peptide backbone, could present an excellent alternative to DNA handles in order to retain addressability in silicified DNA nanostructures. As TMAPS and the peptide backbone in PNA would not be able to electrostatically associate, PNA should remain unsilicified and hence remain available for post-silicification hybridization. Initial studies using a three-strand DNA handle:PNA:anti-PNA handle system (see **Figure S6a**) were promising and showed that indeed, PNA remained accessible for hybridization with anti-PNA coated Au NPs (**Figure S6b**). Nevertheless, due to the high cost of PNA, we also explored more sustainable options. Inspired by work by Ding and co-workers^[8e]^, which showed that TMAPS-TEOS precursors accumulated most favorably on double-stranded DNA (dsDNA) compared to the closely packed dsDNA in a DNA origami nanostructure, we hypothesize that ssDNA may show even less accumulation of silica precursors. We assume that ssDNA, possessing comparatively less phosphate groups compared to dsDNA, may consequently attract less TMAPS molecules. Additionally, we hypothesize that due to the significantly shorter persistence length of ssDNA compared to dsDNA (~ 2 nm^[10]^ vs ~35 nm^[11]^), accumulation of TMAPS could also be minimized, resulting in (mostly) unsilicified strands of ssDNA.

### 2.1. Samples silicified in solution

To test if ssDNA indeed remains largely unsilicified and accessible, we used two different DNA origami nanostructures (a 24 helix bundle (24HB) and a four-layer block (4LB)) displaying ssDNA A_15_-handles protruding from the structure. We recently showed that DNA origami undergo strong condensation during silicification as a result of hydrophobic effects and water depletion forces caused by the influx of silica into the origami structure.^[8g]^ However, even at the maximally condensed state with sub-nm external silica deposition, structures displayed impressive thermal stability. Therefore, we here employed the same structures (24HB and 4LB) as reported in our recent publication and silicified these following our previously published protocol, using a rotator (**Figures S7,8**).^[8g]^ After ~4 h the reaction was stopped, resulting in structures with sub-nm silica deposition that displayed increased stability upon exposure to DNase I (see **Figure S9**). Silicified structures were then incubated with Cy5-labelled T_19_-anti-handles and analyzed by gel electrophoresis. Silicified DNA structures entered the agarose gel and showed similar electrophoretic mobilities to the bare structures (**Figure 1** and **Figure S10**). This is not surprising, since silica deposition in the maximally condensed state is sub-nm, yet the condensation effect^[8g]^ is not drastic enough to influence the electrophoretic mobility significantly. A fluorescent band in the Cy5 fluorescence channel can be clearly observed for the 24HB and the 4LB for both bare and silicified structures displaying the A15-handle, indicating that hybridization to the Cy5-labelled T_19_-anti-handle had been successful. To confirm that this signal is due to specific hybridization rather than non-specific interactions between the Cy5 anti-handle and the silica, we also tested the same origamis without the A_15_-handle.

**Figure 1.**
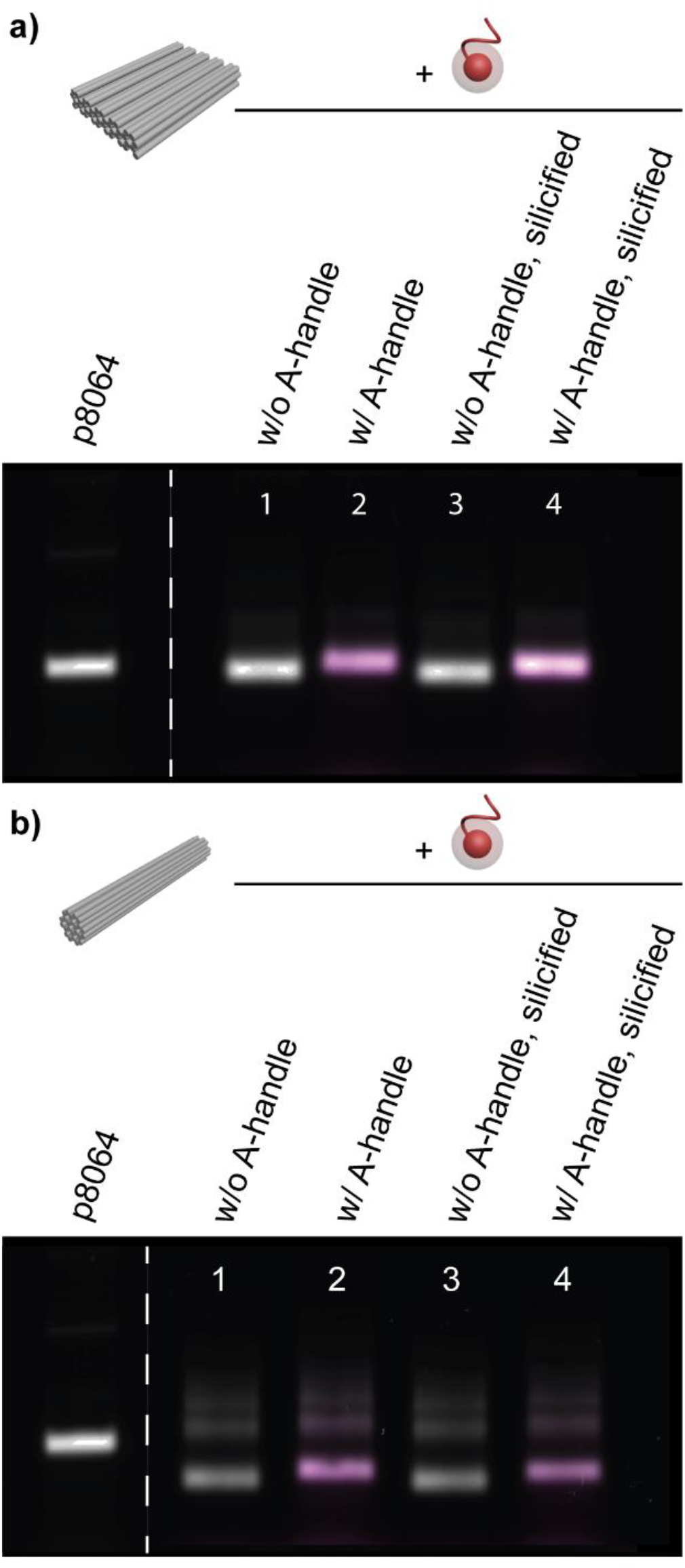
Agarose gel electrophoresis of the 4LB (a) and 24HB (b) before (lanes 1 and 2) and after silicification (lanes 3 and 4) and addition of the Cy5-anti-handle.

As can been seen from **Figure 1** even though structures were incubated with the Cy5 anti-handle, no fluorescent band could be observed. This suggests that a) Cy5 anti-handles successfully hybridized to the ssDNA handles on the origami, b) ssDNA handles remain accessible for hybridization and therefore must be (mostly) unsilicified, c) there is no unspecific interaction between silicified structures and the Cy5 oligonucleotide.

However, as these solution silicified structures only display sub-nm silica deposition as previously established^[8g]^, it could be argued that the retained addressability is not surprising and does not necessarily show that ssDNA handles remain accessible if structures are coated with a thick silica layer. As reported by Liu et *al*. structures silicified on a surface generally display silica layers of several nm thickness.^[8a]^ We therefore next studied the accessibility of ssDNA handles on DNA origami structures immobilized and silicified on a surface.^[8a]^

### 2.2. Samples silicified on surface

Instead of a simple hybridization experiment as carried out for samples silicified in solution, DNA nanostructures immobilized and silicified of on glass surfaces excellently lend themselves for fluorescence imaging studies where the addressability of each DNA origami nanostructure can be assessed on a single particle level. Here we employed DNA-PAINT super-resolution imaging to study the addressability of silicified 12 helix bundle (12HB) DNA nanostructures previously used as super-resolution imaging standards.^[12]^ In DNA PAINT ssDNA docking sites are presented on the molecule of interest (in our case a silicified DNA nanostructure). Short fluorescently-labelled imager strands, complementary to the docking site, then transiently bind from solution allowing for sub-nm localization precision.^[13]^ To investigate whether a short ssDNA handle protruding from a DNA origami would still be accessible if the silica shell measured several nm in thickness, we designed a 12HB containing 8 nt long DNA PAINT docking sites resulting in a distance from the DNA origami surface of only 0 – ~5,4 nm, assuming a length of ~0,67 nm/base in ssDNA^[10b]^ (**Figure 2a**). Initially, to quantify the silica shell thickness on the 12HB, structures were immobilized on mica surfaces and silicified for 4 d. Analysis by atomic force microscopy (AFM) revealed a homogenous height increase of ~5-6 nm (bare vs. silicified origami, **Figure 2b,c** and **Figure S11**). In contrast to the DNA nanostructures silicified in solution with sub-nm external silica deposition, the here observed 5-6 nm thick silica coating on the immobilized 12HB is similar in thickness to the length of the 8 nt DNA PAINT docking site, if fully stretched out. This could have a significantly detrimental effect on the accessibility of the ssDNA docking site. To investigate if such a short docking site would remain accessible at all after silicification, we proceeded with DNA PAINT imaging studies. For this, 12HB DNA origami structures were immobilized on BSA-biotin-streptavidin coated glass coverslips via biotinylated DNA staple strands and silicified for 4 d as before.

**Figure 2.**
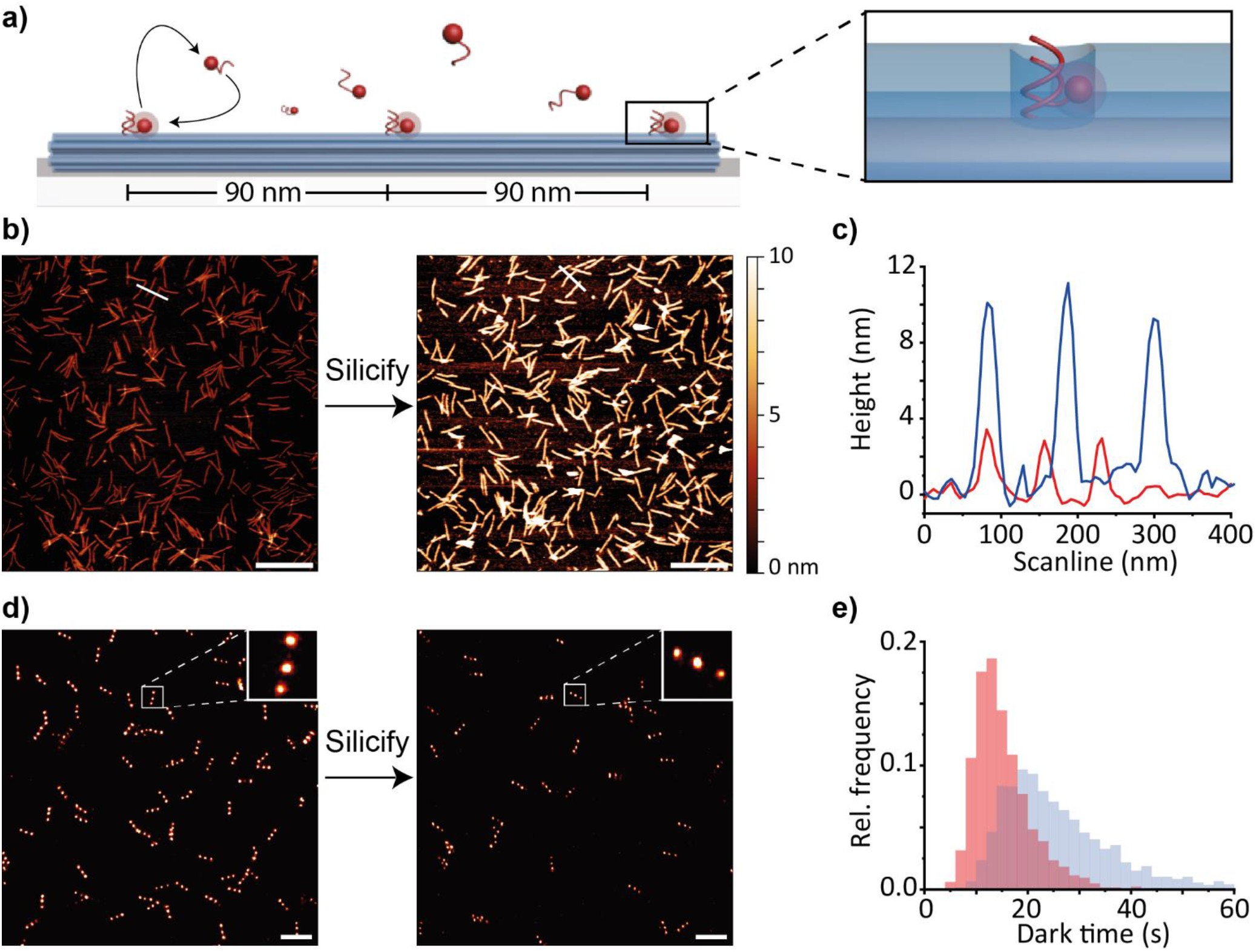
Investigation of DNA origami nanostructures immobilized and silicified on a surface a) Schematic illustration of 12HB DNA origami nanostructure used for DNA-PAINT measurements and illustration of the docking site placement. Inset shows the unsilicified docking site surrounded by silica (blue); b) AFM images of 12HB immobilized on a mica surface before (left panel) and after (right panel) silicification (scale bar: 500 nm); c) height profile of silicified (blue) and bare (red) 12HB nanostructures obtained from AFM images (white lines in b) indicate the line scan); d) super-resolution DNA-PAINT images of 12HB nanostructures before (left panel) and after (right panel) silicification using an Atto655-labelled imager strand (the expected triple spot pattern is shown in the zoomed in images in the insets). Scale bars are 500 nm; e) Extracted distributions of spot integrated dark times for bare (red) and silicified (blue) 12HB nanostructures.

A total of 18 docking sites, arranged in three positions (6 per position to increase binding probability and spot brightness) were incorporated on the 12HB. This three-position arrangement with diffraction limited inter-spot distances of ~90 nm led to a triple spot pattern in a super-resolution DNA-PAINT imaging experiment on the bare 12HB as expected (**Figures 2d**, **S12a**, left panels). Surprisingly, even the silicified 12HB nanostructures displayed well-resolved triple spot patterns (**Figures 2d**, **S12a**, right panel) indicating that the docking site remained unsilicified and still accessible to fluorescently labelled 8 nt long imager strand despite the several nm thick silica layer on the origami.

To gain more information about the accessibility of ssDNA PAINT docking sites we further analyzed the single-molecule binding kinetics obtained from single 12HB nanostructures and extracted the average dark times for each individual labelling spot on the nanostructure (**Figure 2e**). The dark time gives information on the time required for an imager strand to diffuse and hybridize to a docking site. It thus allows to indirectly probe the local accessibility of the docking sites (given that the dissociation time of the 8 nt imager strand occurs at a much faster time scale, i.e., hundreds of ms). As illustrated in **Figure 2e**, the dark time distribution for silicified 12HB structures was significantly broader compared to that of bare 12HB structures. Additionally, we observed a slight shift of the dark time distribution to longer time scales (from 15.2 ± 5.7 s mean dark time for bare 12HB to 26.9 ± 15.4 s for silicified 12HB). This suggests that the change in local microenvironment around the partially embedded docking site as a result of silicification, resulted in substantially slowed down diffusion kinetics of incoming imager strands. Nevertheless, taken together, our findings strongly suggest that ssDNA remains unsilicified and may thus result in the formation of a small pore within the silica shell, around the ssDNA.

As the here employed 12HBs are commonly used as fluorescent nano rulers, whose quality and overall lifetime could be greatly improved with increased stability, we next assessed the stability of the silicified structures and their handles in degrading buffer conditions. For this we incubated both silicified and non-silicified 12HB nanostructures, immobilized on a glass coverslip (as described above), in 1× TAE buffer for 2 h. It was previously reported that DNA origami nanostructures displayed low stability in the absence of or at low concentrations of Mg^2+^ ions in the presence of EDTA.^[14]^ Therefore, it was not surprising that bare 12HB structures no longer displayed the representative triple spot pattern in the DNA-PAINT localization image. Instead, we observed most of the localizations clustered in one spot, indicating structural collapse (**Figure S12c**, left panel). In contrast, silicified 12HB nanostructures remained intact and the expected triple-spot pattern was still observable, confirming successful silicification and improved stability of the 12HB sample (**Figure S12c**, right panel). This also illustrates the potential of the silicification approach with retained addressability to extend the utility of functional DNA nanostructures to applications typically limited by the stability of DNA origami in harsh handling conditions, such as low/no salt containing buffers or even non-aqueous solutions, which could be especially important for materials science applications.^[15]^ For this it would also be interesting to obtain dynamic DNA nanostructures with the mechanical resilience of an inorganic material, but the structural shape-changing flexibility of a DNA nanostructure. We therefore next sought to test if stretches of ssDNA inside a DNA origami structure could also remain accessible for hybridization, allowing for the formation of flexible, shape-changing structures.

### 2.3. Dynamic DNA origami

To obtain a dynamic, flexible DNA origami with shape-changing properties, we omitted a set of staples from the middle of an 18HB (see **Figure 3a and S13**) leaving only the scaffold to connect the two halves of the structure. As expected, this resulted in a flexible structure, appearing significantly bent upon deposition on a TEM grid (**Figure 3b left panel**). Observed bending angles ranged from 15 to 180° with the majority of structures displaying bending angles between 120 and 150° (**Figure 3d left panel**). However, a subsequent addition of the missing middle staples (hereafter referred to as “straightening staples”) and incubation at 36 °C resulted in a distinct shift in bending angles (**Figure 3d,** left panel) with a majority of structures straightening out as evidenced by TEM analysis (**Figure 3b,** right panel). A similar effect had previously also been observed for 12HB structures.^[16]^ After successfully confirming that bent 18HBs can be straightened out after addition of the straightening staples, we next tested if this was still possible for silicified structures. Bent 18HB were hence silicified using the solution approach.^[8g]^ TEM analysis revealed that silicified structures also appeared bent as expected, confirming their retained flexibility (**Figure 3c,** left panel). Structures showed a similar trend in observed bending angles with most structures displaying bending angles between 135 – 180° (**Figure 3d,** right panel). However, we also observed that silicified structures on average tended to display slightly larger bending angles, presumably due to the increased stiffness inferred by the silica. In order to test if the ss scaffold sections in the middle of the 18HB were still accessible for hybridization after silicification, we added the straightening staples and incubated the mixture at 36°C as described above. Analysis by TEM revealed a clear shift towards a 180° angle (i.e. straight structures), indicating that ss scaffold segments within a DNA origami also remained largely unsilicified and accessible for further hybridization (**Figure 3c**, right panel). However, the amount of fully straight structures was slightly less for silicified samples compared to bare ones. We hypothesize that this is on the one hand due to increased flexibility of dsDNA compared to inorganic silica. On the other hand we assume partially obstructed diffusion of the straightening staples into the silicified halves of the 18HB, similar to the slowed-down diffusion kinetics observed for the imager strands in the DNA PAINT imaging experiments. Nevertheless, our study strongly suggests that the formation of dynamic, flexible, shape-changeable silica-DNA hybrid nanostructures is indeed possible, opening up new possibilities for applications in biosensing, materials science or even nano robotics.

**Figure 3.**
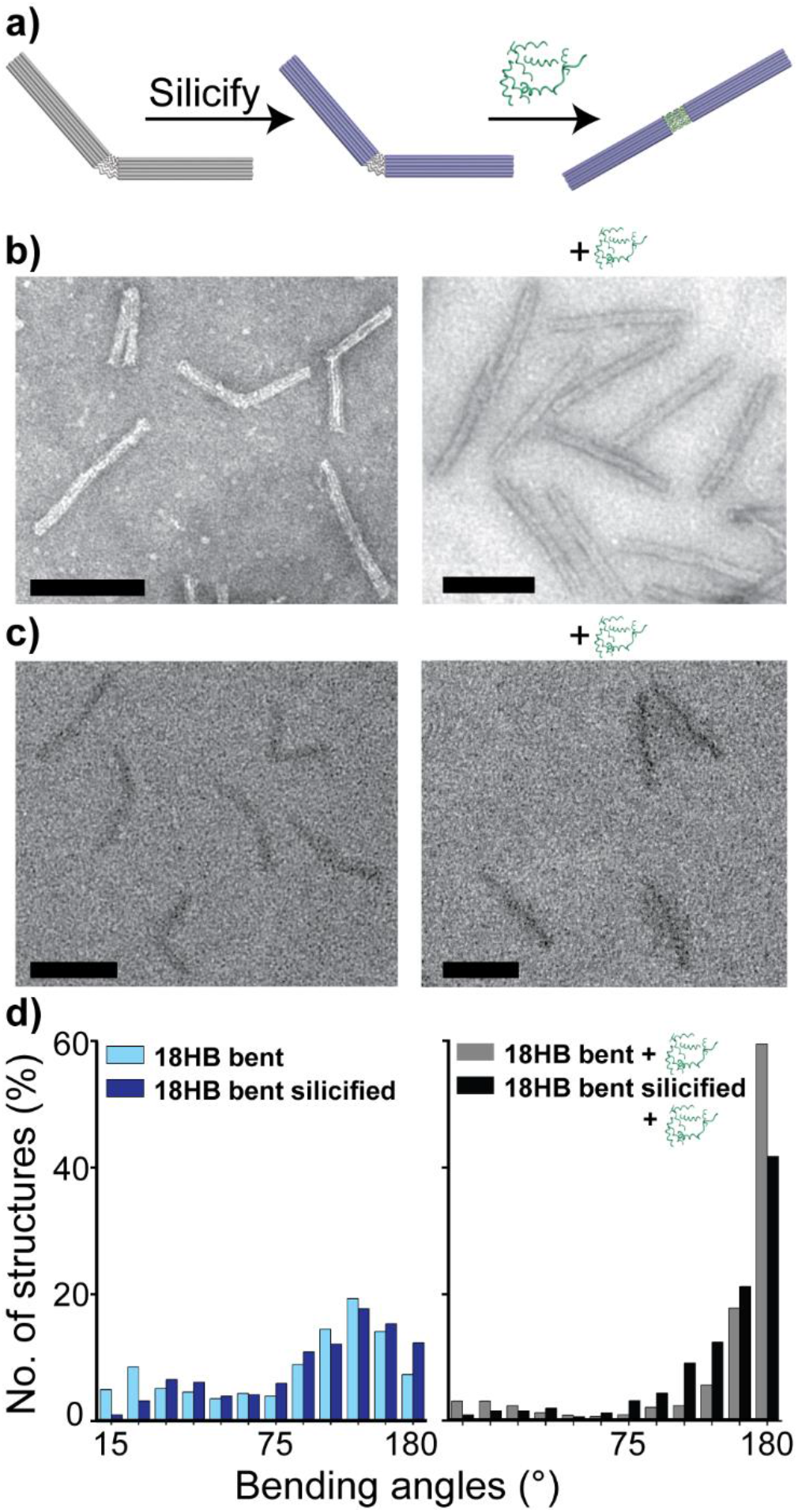
(a) Schematic illustration of a bent 18HB with missing middle staples. After silicification (blue structure) and subsequent addition of the corresponding straightening staples (green), structures straighten out. (b) Bare and (c) silicified 18HB before and after addition of the corresponding straightening staples. Bare structures were stained with uranyl formate, while silicified structures were not stained. Scale bars are 100 nm. (d) histograms of bending angle before (left) and after addition of straightening staples (right). More than 480 structures were analyzed for each condition. (Angle distributions were collated in 15° bins).

### 2.4 DNA origami-silica crystals from silicified monomers

Finally, we aimed to demonstrate the power of silicification with retained addressability for materials science applications. DNA-programmable nanomaterial crystals have gained a lot of attention in recent years. Employed materials range from quantum dots to proteins, plasmonic NPs and DNA origami.^[4]^ However, only very few DNA origami designs so far showed the ability to form single crystals.^[4]^ A prominent type of DNA origami showing excellent crystal formation abilities are polyhedra. Generally, DNA-programmable lattices must be silicified after their formation in order to allow for their analysis in a dry state, avoiding structural collapse.^[8b, 17]^ Here we show that crystals based on sticky ended hybridization interactions of monomers can also be formed from *pre-silicified* monomers (**Figure 4**). We designed an octahedral DNA origami monomer - inspired by the work of Wang et *al*.^[17b]^ - made up of twelve 6HB edges. Four sticky end sequences at each vertex allow for the formation of cubic microcrystals.

**Figure 4.**
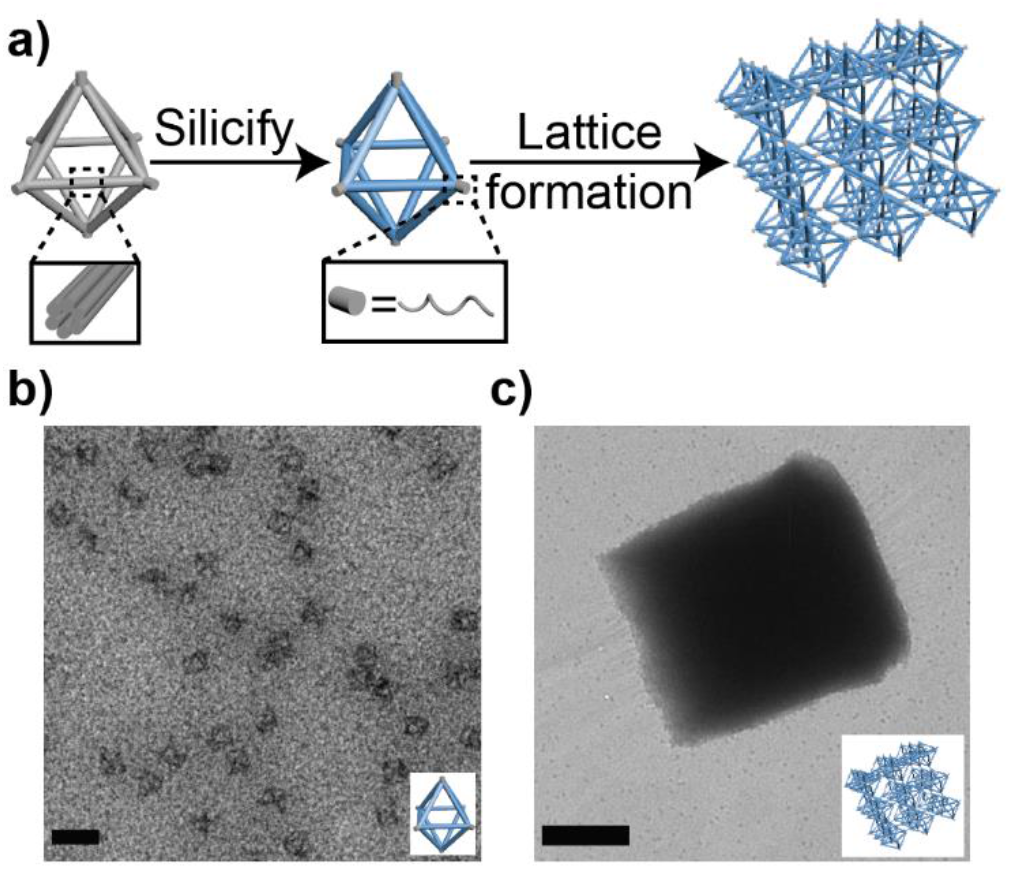
(a) Schematic illustration of DNA origami crystal formation. Octahedral DNA origami monomers made up of 6HB struts were silicified in solution and subsequently formed 3D crystals via sticky end hybridization. (b) TEM images of silicified octahedra and (c) corresponding TEM image of a silicified DNA origami crystal, both unstained. Scale bars are 100 nm (a) and 1 μm (b).

Octahedra were silicified in solution as described before. However, due to their small size and delicate nature, high losses were observed during ultrafiltration, resulting in average obtainable concentrations of only ~50 nM. Therefore, structures were silicified at this comparatively lower concentration, adjusting the concentrations of TMAPS and TEOS accordingly and still maintaining rotation during silicification. Analysis by TEM revealed that silicified octahedra were well visible without additional staining with uranyl formate. Structures also appeared less deformed and more 3D in nature compared to bare, stained structures (see **Figure 4b** and **Figure S14**). This is most likely due to the increased stiffness inferred by the silica. To further test the stability of the silicified octahedra, we exposed them to 60 °C heat for 30 min. Subsequent analysis by TEM confirmed that structures remained largely intact (**Figure S14**), suggesting that silicification even at lower concentrations inferred substantial thermal stability. Having established the successful silicification of the octahedra, we next turned our attention to the crystal formation. Based on our previous findings, the ssDNA sticky ends should remain unsilicified and hence allow for hybridization and subsequent lattice formation. Silicified monomers were therefore incubated and exposed to a temperature ramp (see **materials and methods**) followed by analysis by TEM. As can be seen in **Figure 4c** the silicified monomers were still capable of forming several micrometer-sized cubic single crystals. Compared to bare DNA origami crystals, crystals formed from silicified monomers appeared more 3D in nature (see **Figure S15**), suggesting that the stiffness inferred by the silicification allows crystals to better retain their 3D shape in a dry state. To date it has been very challenging to form open-channel 3D crystal lattices from inorganic materials^[18]^, however, our findings show that this can be easily achieved using DNA origami-templated silica nanostructures.

## 3. Conclusion

Many attempts to explore potential real-life applications of DNA origami have faced the trouble of its inherent instability in non-aqueous conditions or those commonly met within biological environments. Silicification of DNA origami has helped to overcome the stability bottleneck. However, it was thus far believed that silicification renders the resulting nanostructures no longer site-specifically modifiable with other functional molecules through hybridization. Here we were able to show that this is not the case. In summary, we have demonstrated that ssDNA handles as well as ssDNA scaffold segments remain unsilicified both for solution and “on surface” silicification approaches independent of the degree of silica coating. The silica nanostructures are precisely templated by the DNA origami, while the most attractive feature of DNA nanostructures – complete and accurate addressability – can be retained. This brings an interesting and important new feature to silica nanostructures. It allows for tight control over the conjugation of functional molecules and materials (e.g. fluorophores, NPs, quantum dots, proteins), both spatially and numerically. The silica-DNA hybrid crystals formed here will also allow to strategically and specifically place functional molecules inside the inorganic crystal and could even allow for a controlled assembly and disassembly processes without affecting the monomers. Our finding of a fully site specifically addressable inorganic nanostructure with complete control over size and shape opens up a new era for silica nanostructures by combining the robustness of an inorganic material with the full power of DNA self-assembly and complete and accurate addressability, harnessing the excellent properties of both materials. This will allow for and inspire new and exciting applications ranging from biomedicine and catalysis to materials science.

## Data availability

All data supporting the key findings of this study are available within the main text and supplementary information files.

## Author contributions

A.H-J conceived the idea. L.M.W., A.V.B and M.S. fabricated samples and carried out experiments and data analysis. M.S. carried out DNA PAINT and AFM imaging. A.H-J and V.G. supervised the study. A.H-J and V.G. wrote the manuscript. All authors discussed and edited the manuscript and gave approval to the final version of the manuscript.

## Acknowledgements

We acknowledge financial support from the German Research Foundation (DFG) through SFB1032 (Nanoagents) project A06 and the Emmy Noether program (project no. 427981116) (A.H-J). V.G. gratefully acknowledges financial support from the DFG (grant number GL 1079/1-1, project number 503042693). This work was also financially supported by the Center for Nanoscience (CeNS) through a collaborative research grant to A.H-J and V.G. We thank Marianne Braun and Ursula Weber for assistance with TEM imaging.

## 4 Supporting Information

### Note S1: DNA origami designs

DNA origami structures were designed using the caDNAno software^[1]^ (design schematics in Figures S1–S5).

**Figure S1:**
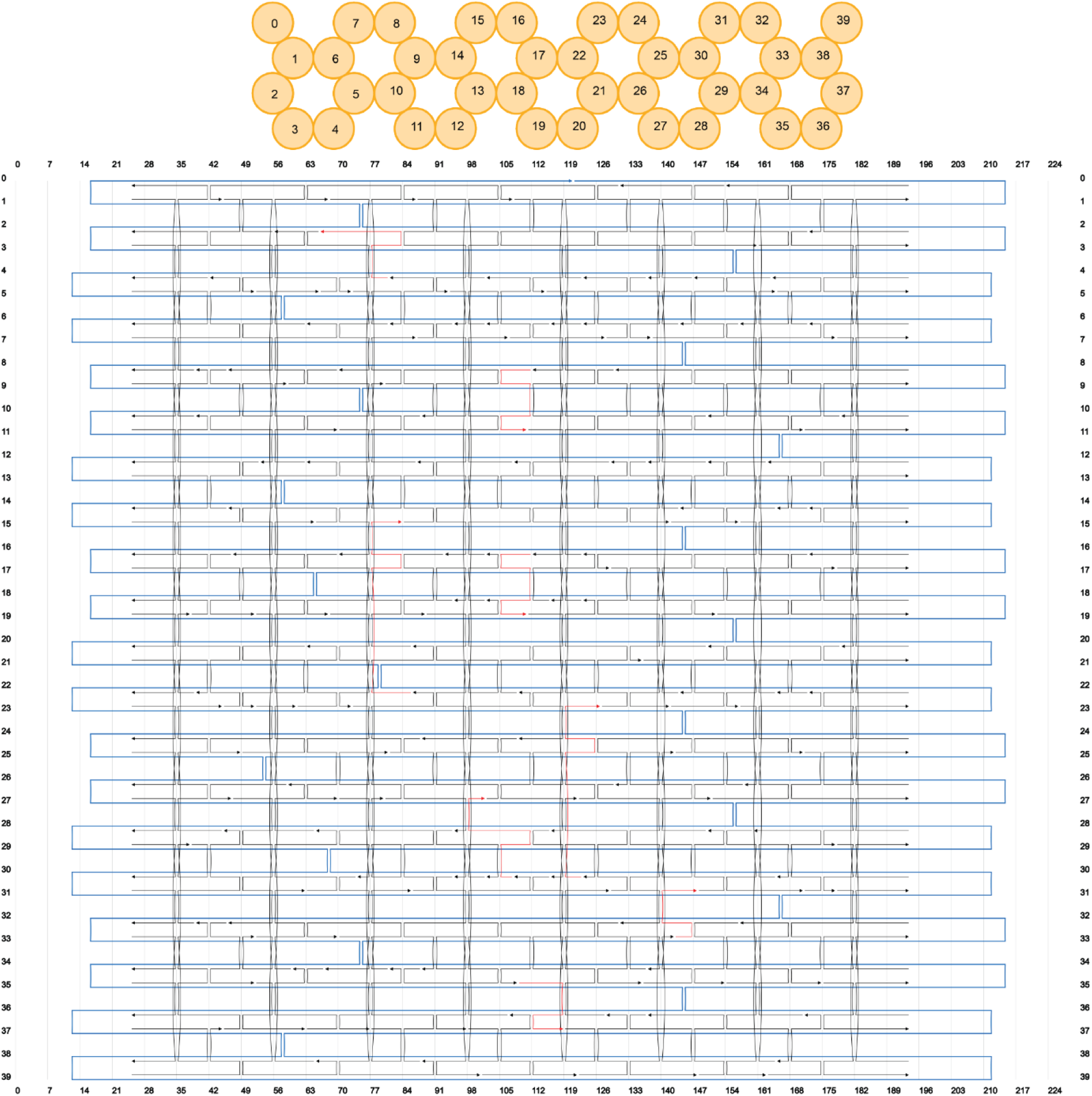
CaDNAno scaffold (blue, p8064) and staple paths (black) of the four-layer block (4LB) structure. The staples marked in red are A_15_-extended handles (extension at 5’ end).

**Figure S2:**
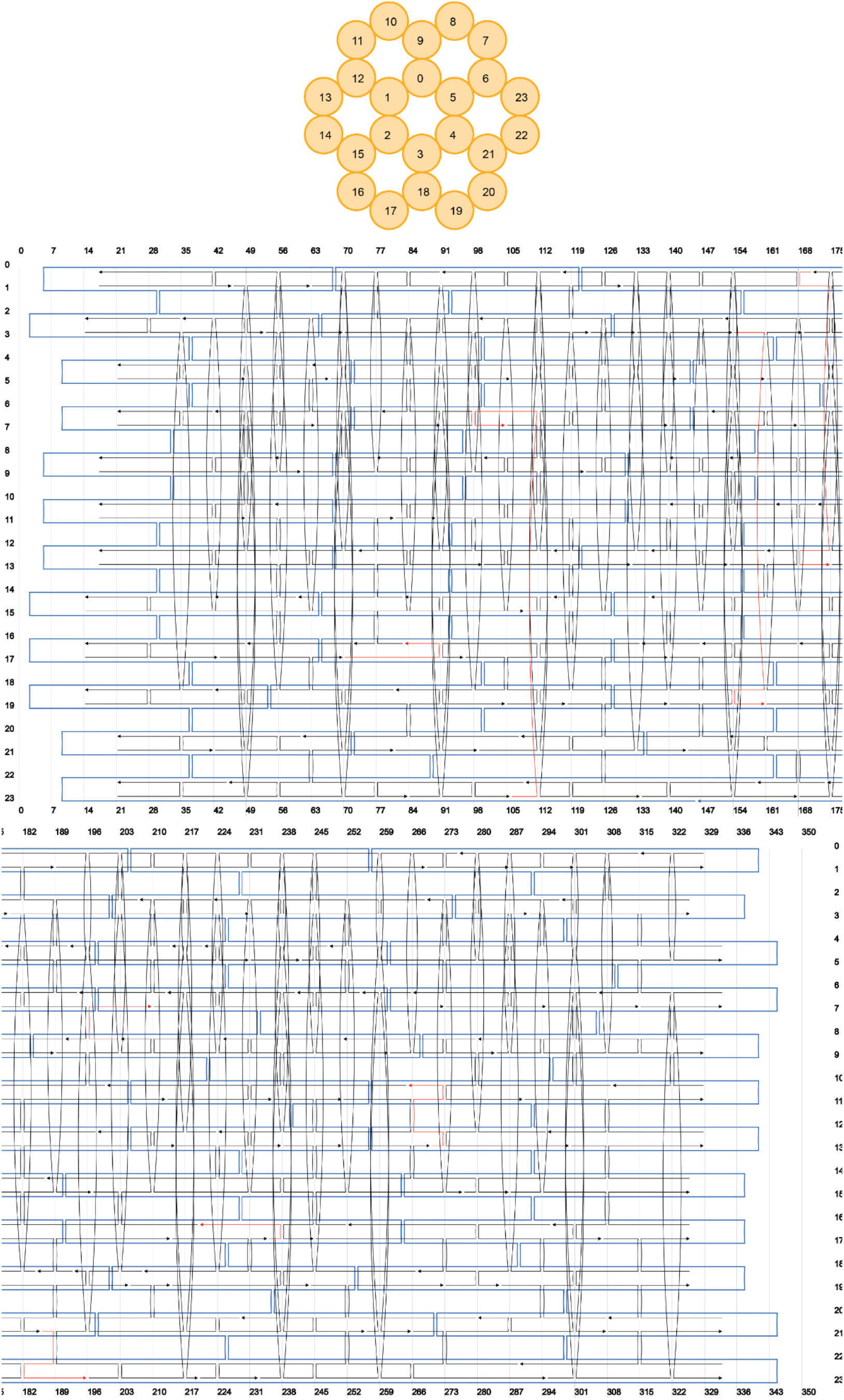
CaDNAno scaffold (blue, p8064) and staple paths (black) of the 24 helix bundle (24HB) structure. The staples marked in red are A15-extended handles (extension at 5’ end).

**Figure S3:**
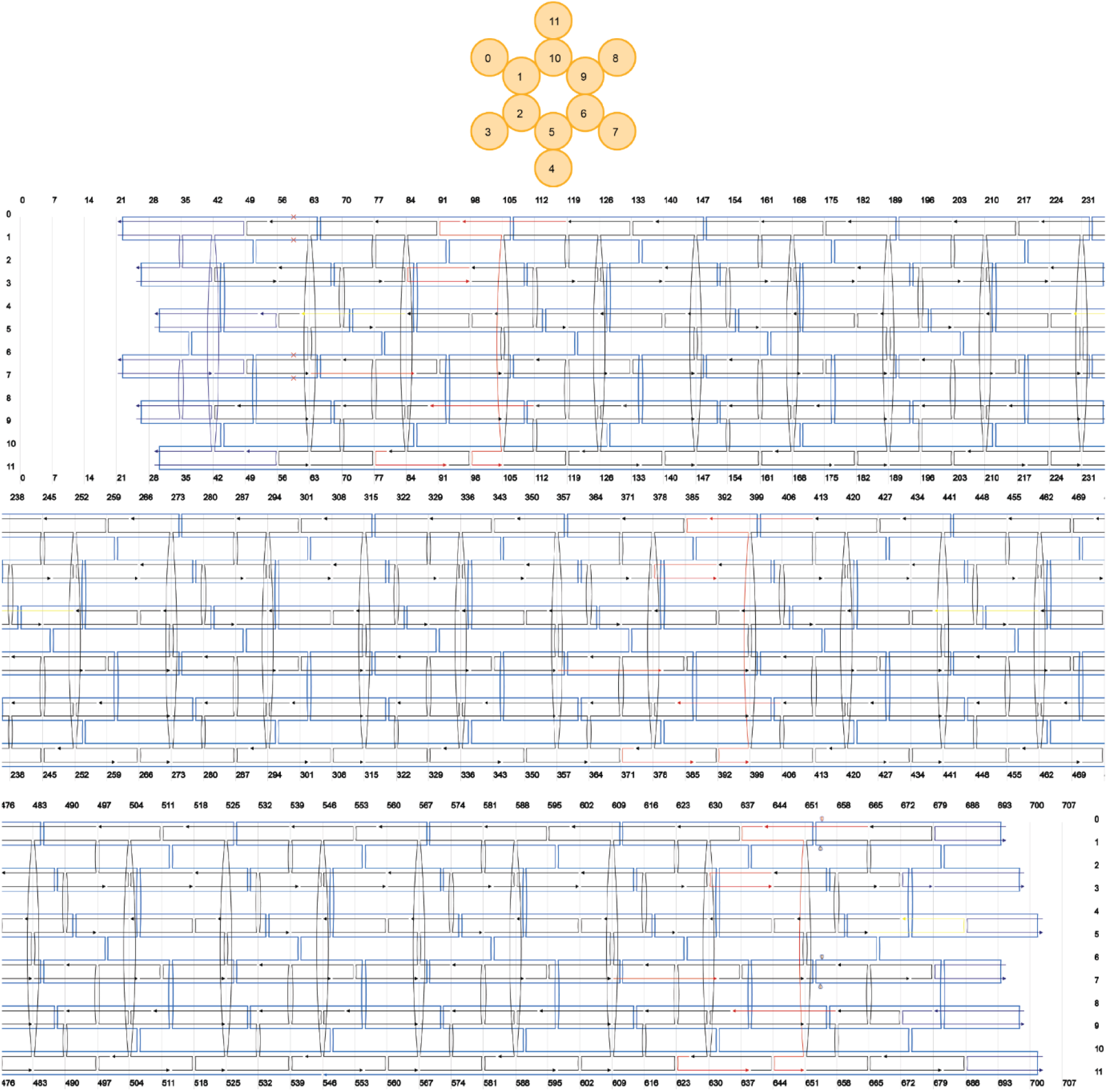
CaDNAno scaffold (blue, p8064) and staple paths (black) of the 12 helix bundle (12HB) structure.^[2]^ The red staples represent DNA PAINT staples with docking sites of a 8 nt binding sequence on the 3’-end. Yellow staples denote biotinylated staples for surface immobilization.

**Figure S4:**
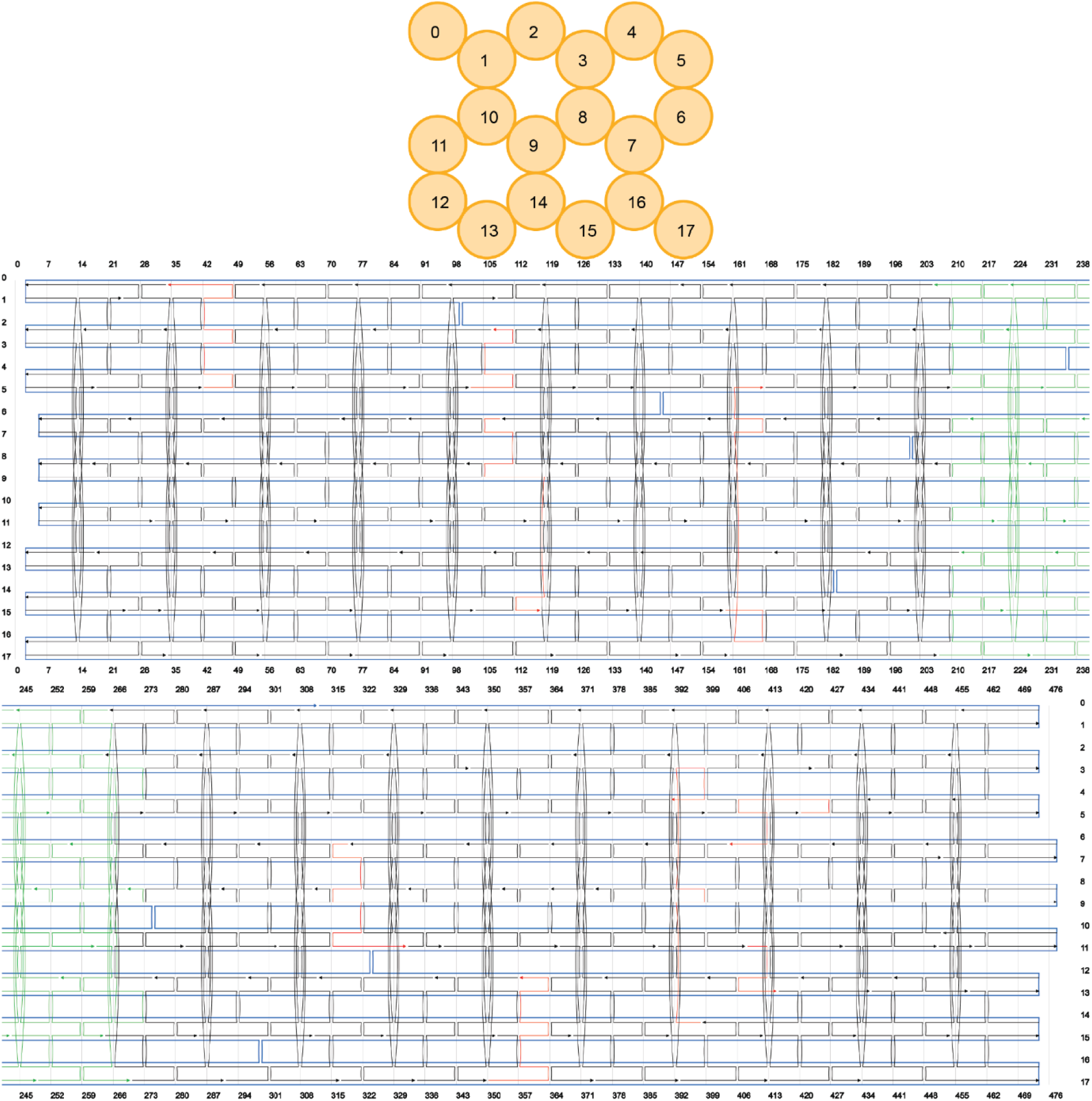
CaDNAno scaffold (blue, p8634) and staple paths (black) of the 18 helix bundle (18HB) structure. The staples marked in red are A_15_-extended handles (extension at 5’ end). The staples marked in green were omitted for the bent 18HB structure.

**Figure S5:**
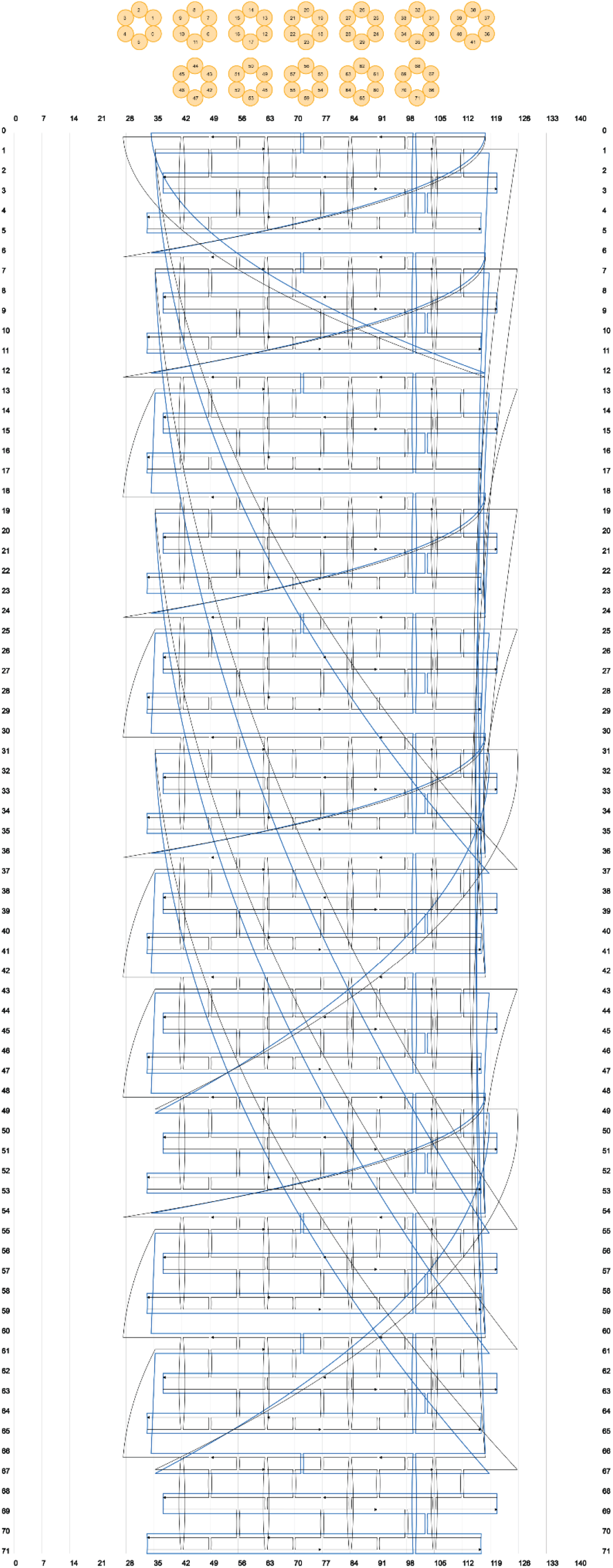
CaDNAno scaffold (blue, p7249) and staple paths (black) of the octahedron. The design was created using TALOS^[3]^ and caDNAno.

### Note S2: Materials and methods

#### Chemicals

Unless stated otherwise, all chemicals were used as received. Tetraethylorthosilicate 98 % (TEOS) and MgCl_2_ 98 % were obtained from Sigma Aldrich, Trimethyl(3-(trimethoxysilyl) propyl)ammonium chloride (50 % in methanol) (TMAPS) was obtained from TCI America. Oligonucleotides were obtained from Eurofins Genomics or IDT. DNase I was obtained from Roche. 10×DNase I buffer was obtained from New England Biolabs.

#### Folding of DNA origami structures

All DNA origami structures used here were designed using the CaDNAno software (design schematics in note S1).

##### 4 Layer Block (4LB)

The 4LB was folded using 10 nM of the scaffold p8064, 100 nM of each staple strand in buffer containing 5 mM Tris, 1 mM EDTA (pH = 8) and 20 mM MgCl_2_. The mixture was heated to 65 °C and held at this temperature for 30 min, then cooled down to 4 °C over a period of 16 hours. All additional handle staples were incorporated during folding (see **Figure S1** for handle positions and **note S12** for specific sequences).

##### 24 Helix Bundle (24HB)

The 24HB was folded using 10 nM of the scaffold p8064, 100 nM of each staple strand in buffer containing 5 mM Tris, 1 mM EDTA (pH = 8) and 18 mM MgCl_2_. The mixture was heated to 65 °C and held at this temperature for 30 min, then cooled down to 4 °C over a period of 16 hours. All additional handle staples were incorporated during folding (See **Figure S2** for handle positions and **note S12** for specific sequences).

##### 12 Helix Bundle (12HB)

The 12HB DNA origami was folded using 20 nM of the scaffold p8064, 200 nM of each unmodified staple strand and 600 nM of each modified staple strand (biotinylated and DNA PAINT staple strands) in buffer containing 50 mM Tris, 20 mM acetic acid, 1 mM EDTA (pH = 8) and 16 mM MgCl_2_. The mixture was heated to 65 °C and then cooled down to 4 °C over a period of 25 hours with a non-linear thermal annealing ramp adapted from ref.^[4]^. All additional handle staples were incorporated during folding (see **Figure S3** for handle positions and **note S12** for specific sequences).

##### 18 Helix bundle (18HB)

The 18HB DNA origami structure was folded using 10 nM of the scaffold p8634, 100 nM of each staple strand in buffer containing 5 mM Tris, 1 mM EDTA (pH = 8) and 18 mM MgCl_2_. The mixture was heated to 65 °C and held at this temperature for 30 min, then slowly cooled down to 4 °C over a period of 16 hours. To achieve the bent structure in the 18HB, 25 staples from the middle of the 18HB were not included in the folding procedure (see **Figure S5** and **note S12** for omitted staples). This results in a 18HB with a single-stranded (scaffold-only) part in its middle where the two fully folded parts can move independently from each other, giving the structure the appearance of being bent. The missing staples were added in a 10-fold molar excess after silicification to straighten the 18HB back out and the mixture was kept at 36°C for 16 hours to guarantee incorporation.

#### Purification of DNA origami structures

All folded DNA origami structures were purified from excess staple strands via ultrafiltration (Amicon filter units, 100 kDa). Briefly, the folding mixture (~2 mL) was divided over 2 Amicon Ultra filters (0.5 mL, 100 K, Millipore, USA) and each centrifuged at 8,000 rcf for 8 min. The centrifugal steps were repeated up to 10 times (until no staples were detectable in the flow through) with fresh buffer (1×TAE, 3 - 11 mM MgCl_2_) added in every step.

The successful folding of structures was confirmed by TEM or AFM analysis. DNA origami solutions were stored at −20 °C until further use.

#### DNA origami silicification

##### Silicification in solution

Adapting our previously established protocol^[5]^, unless stated otherwise, all DNA origami solutions used had a concentration of 200 nM and were dispersed in 1× TAE buffer containing 3 mM MgCl2 (50 μL total reaction volume). The sample was placed on a thermo shaker and the first silica precursor TMAPS (TCI, diluted 1:19 in methanol) was added to the sample in 5-fold molar excess to the number of nucleobases. After one minute of shaking at 300 rpm at 21°C, TEOS (Sigma Aldrich, diluted 1:9 in methanol) in 12.5-fold molar excess to the number of nucleobases was added to the solution. The sample was then transferred to a tube revolver rotator (Thermo Scientific) and rotated at 40 rpm at 21 °C for 4 h to reach the “maximally condensed state”^[5]^. Following this, the silicified sample was purified once via ultrafiltration (Amicon filter, 30kDa). For this purpose, the silicified sample was loaded into a pre-washed filter unit and 400 μL of fresh MilliQ water were added. The filter was then centrifuged for 4 min at 8000 rpm. Finally, the DNA origami were eluted by inverting the filter, placing it in a new tube and centrifuging the new tube for 3 min at 5000 rpm.

##### Silicification on surface

For surface silicification, the well-established literature protocol by Fan and co-workers was adapted.^[6]^ DNA origami samples were either immobilized on glass slides (see “Glass surface preparation”) or on mica. Initially, a precursor solution was prepared by adding 1 mL of 1× TAE–Mg^2+^ buffer (40 mM Tris, 2mM EDTA-Na_2_, 12.5 mM MgAc_2_, pH=8.0) to a 10 mL glass bottle with a suitably-size magnet and then slowly adding 20 μL of TMAPS (50% (wt/wt) in methanol). This solution was then stirred vigorously for 20 min at room temperature. After that, 20 μL of TEOS were slowly added and the resulting solution was again stirred for 20 min at room temperature. Finally, 400 μL of the precursor solution were immediately transferred to the glass slide containing the immobilized DNA origami. Alternatively mica slides containing adsorbed DNA origami were placed on top of a large precursor droplet on a small petri dish as described in detail in the literature^[6b]^. The glass slide or petri dish was closed airtight and was then gently shaken for 60 min at 40 rpm at room temperature, the samples were left undisturbed for up to 5 days. Afterwards the samples were washed once with 400 μL 80% ethanol and once with 400 μL MilliQ water. Then the samples were stored with a sufficient amount of MilliQ to prevent drying and the samples were sealed airtight again until analysis.

#### Assessing handle accessibility

To determine if ssDNA handles were still accessible for hybridization after silicification, structures were designed to display protruding A_15_-handles (see **note S1** for design information). After purification (and optional silicification) complementary Cy5-labelled T_19_-anti-handles (biomers.net) were added to the origami solution in a 10-fold molar excess and the sample was kept at 36 °C for 16 h prior to analysis by agarose gel electrophoresis (AGE).

#### DNase stability tests

DNase stability tests were conducted according to established literature protocols.^[7]^ Briefly, (silicified) DNA origami (10 nM, 45 μL in 1× TAE buffer containing 3 mM MgCl_2_) were mixed with 10× DNase I buffer (5 μL, NEB) and then split evenly into five 1.5 mL tubes, and added to a thermo mixer (Eppendorf) at 37 °C. DNase I (1 μL, 0.1 U/μL, NEB) was then added consequentially to one tube each to react for predetermined amounts of time (10 min, 20 min, 30 min and 60 min). As a reference, 1 μL nuclease free water instead of DNase I was added to the last tube (0 min reaction time). Reactions were subsequently quenched by putting the tubes on ice. Samples were then analyzed by TEM.

#### Agarose Gel Electrophoresis (AGE)

DNA origami samples (10 μL, diluted to 10 nM in 1× TAE buffer containing 3 mM MgCl_2_) were mixed with loading buffer containing orange G and Ficoll, and loaded onto a 0.7% agarose gel (1× TAE, 11 mM MgCl_2_), which was stained with 0.01% SYBRSafe. Gels were run on ice for 90 min at 75 V (running buffer: 1× TAE with 11 mM MgCl_2_). Gel imaging was subsequently carried out using the Typhoon FLA-9000 (GE Healthcare).

#### Transmission electron microscopy (TEM)

DNA origami sample (10 nM, 10 μL) was applied to a plasma-cleaned, carbon-coated copper grid that had been plasma-cleaned for 30 seconds. Bare DNA origami samples were incubated for 90 s and the remaining solution was removed with filter paper. Afterwards samples were stained with 2% uranyl formate (5 μL) solution for 30 seconds. Silicified DNA origami samples were incubated on the grid for 10 minutes, before the remaining solution was removed using a filter paper. The grid was then washed once with MilliQ water and dried in air before imaging. Images were obtained on a Jeol-JEM-1230 TEM operating at an acceleration voltage of 80kV. Images were subsequently analyzed using the ImageJ software.

#### Atomic force microscopy (AFM)

AFM scans in aqueous solution (AFM buffer = 40 mM Tris, 2 mM EDTA, 12.5 mM Mg(OAc)_2_·4 H_2_O) were realized on a NanoWizard^®^ 3 ultra AFM (JPK Instruments AG). For sample immobilization, a freshly cleaved mica surface (Quality V1, Plano GmbH) was incubated with 10 mM solution of NiCl_2_ for 3 minutes. The mica was washed three times with ultra-pure water to get rid of unbound Ni^2+^ ions and blow-dried with air. The dried mica surface was incubated with 1 nM sample solution for 3 minutes and washed with AFM buffer three times. Measurements were performed in AC mode on a scan area of 3 x 3 μm with a BioLeverMini cantilever (ν_res_ = 110 kHZ air / 25 kHz fluid, k_spring_ = 0.1 N/m, Bruker AFM Probes).

Leveling, background correction and extraction of height histograms of obtained AFM images were realized with the software Gwyddion^[8]^ (version 2.60).

#### Glass surface preparation

For optical microscopy experiments, the DNA origami sample was immobilized on Nunc^™^ LabTek^™^ II chambers (Thermo Fisher, USA). The chambers were first cleaned with 500 μL of 1% HellmanexIII^™^ solution (Sigma Aldrich, USA) overnight and washed thoroughly with water, then three times with 1× PBS buffer. Then the surfaces were passivated with 100 μL BSA-biotin (0.5 mg/mL in PBS, Sigma Aldrich, USA) for 15 min and washed three times with 1 × PBS buffer. The passivated surfaces were incubated with 100 μL streptavidin (0.25 mg mL-1 in PBS, S4762, Sigma Aldrich, USA) for 15 min and washed three times with 1 × PBS buffer. The sample solution with DNA origami featuring several staple strands with biotin modifications on the base was diluted to approximately 200 pM in 2× PBS buffer containing 500 mM NaCl and incubated in the chambers for 5 to 15 minutes. Sufficient surface density was probed with a TIRF microscope.

#### DNA PAINT

DNA PAINT measurements were carried out on a custom-built total internal reflection fluorescence (TIRF) microscope, based on an inverted microscope (IX71, Olympus) placed on an actively stabilized optical table (TS-300, JRS Scientific Instruments) and equipped with a nosepiece (IX2-NPS, Olympus) for drift suppression. The sample was excited at 644 nm with a 150 mW laser (iBeam smart, Toptica Photonics). The laser beam was spectrally cleaned up (Brightline HC 650/13, Semrock), directed over a dichroic mirror (zt 647 rdc, Chroma) and focused on the back focal plane of the objective (UPLXAPO 100×, NA = 1.45, WD = 0.13, Olympus). An additional ×1.6 optical magnification lens was applied to the detection path resulting in an effective pixel size of 101 nm. The fluorescence light was spectrally filtered with an emission filter (ET 700/75, Chroma). Image stacks in TIF format were recorded by an electron multiplying charge-coupled device camera (Ixon X3 DU-897, Andor), which was controlled with the software Micro-Manager 1.4.^[9]^

All DNA PAINT measurements were conducted at ca. 1.8 kW/cm^2^ (measured power before microscope body) at 640 nm in TIRF illumination with an exposure time of 100 ms and 18,000 frames over 30 min.

For imaging, a 2× PBS buffer containing 500 mM NaCl and 0.05% Tween20^®^ (Sigma Aldrich, USA) and an imager concentration of 10 nM was used. The 8 nt imager oligonucleotide was purchased from Eurofins Genomics GmbH (Germany) and consisted of the sequence 5-GGAATGTT-3 with an Atto655 label on the 3’-end.

Acquired DNA PAINT raw data were analyzed using the Picasso software package.^[10]^ The obtained TIF-movies were first analyzed with the “localize” software from Picasso. For fitting the centroid position information of single point spread functions (PSF) of individual imager strands, the MLE (maximum likelihood estimation) analysis was used with a minimal net gradient of 5000 and a box size of 5 px. The fitted localizations were further analyzed with the “render” software from Picasso. x-y-drift correction of the localizations was corrected with the RCC drift correction.

#### DNA origami crystal formation

Octahedral DNA origami monomers were folded using 20 nM of the scaffold p7249 and 100 nM of each staple strand in buffer containing 5 mM Tris, 1 mM EDTA (pH = 8) and 12.5 mM MgCl_2_. Two mixtures containing the different end staples (type A or type B) were prepared. The mixtures were heated to 95 °C and held at this temperature for 1 min, then cooled down to 20 °C over a period of 20 hours.

The folded DNA origami nanostructures were purified via ultrafiltration (Amicon centrifugal filter units, 0.5 ml, 100 kDa cut-off). The folding mixture was loaded into the filter units and each centrifuged at 2,000 rcf for 20 min. The centrifugal steps were repeated 5 times with fresh buffer (1× TAE, 7.5 mM MgCl_2_) added in every step.

Silicification of octahedral DNA origami monomers was carried out similarly to the procedure described above. Here, 50 μL of DNA origami sample were prepared at a concentration of 50 nM in 1× TAE, 7.5 mM MgCl_2_ buffer. After 4 h of silicification on the rotator, the silicified samples were purified from excess silica via one round of ultrafiltration, as described above. Polymerization into crystalline lattices was carried out by mixing the two different types (A and B) of silicified and purified monomers in a 1× TAE buffer containing 20 mM MgCl_2_. The sample was heated to 48 °C for one hour and then very slowly and gradually cooled down to 20 °C (−1 °C per 150 min).

For the TEM grid preparation for the crystals made from silicified monomers, 10 μL of the sample was taken from the bottom of the tube and applied to a grid that had been plasma-cleaned for 30 s. The sample was incubated for 45 to 55 min on the grid followed by removal of the remaining solution using a filter paper. Afterwards, the TEM grid was carefully washed twice with 5 μL of MilliQ water each and then air-dried before imaging.

### Note S3: Retained addressability through PNA handles

Initial studies on retained handle addressability were carried out using a three-strand system where 1LS were designed with three protruding handles, resulting in one binding site (Figure S6a). To the handles a partially complementary PNA handles (Table S1) was hybridized prior to silicification. Due to a lack of charge on the PNA, no electrostatic association with TMAPS should be possible. After silicification, following ref.^[11]^, silicified 1LS were incubated with a 10× excess of 15 nm Au NPs functionalized with thiolated anti-PNA handle (Table S1). Samples were then purified from excess Au NPs by AGE and subsequently analyzed by TEM (Figure S6b), clearly showing silicified 1LS conjugated to an Au NP, suggesting that PNA remained addressable after silicification as hypothesized.

**Figure S6.**
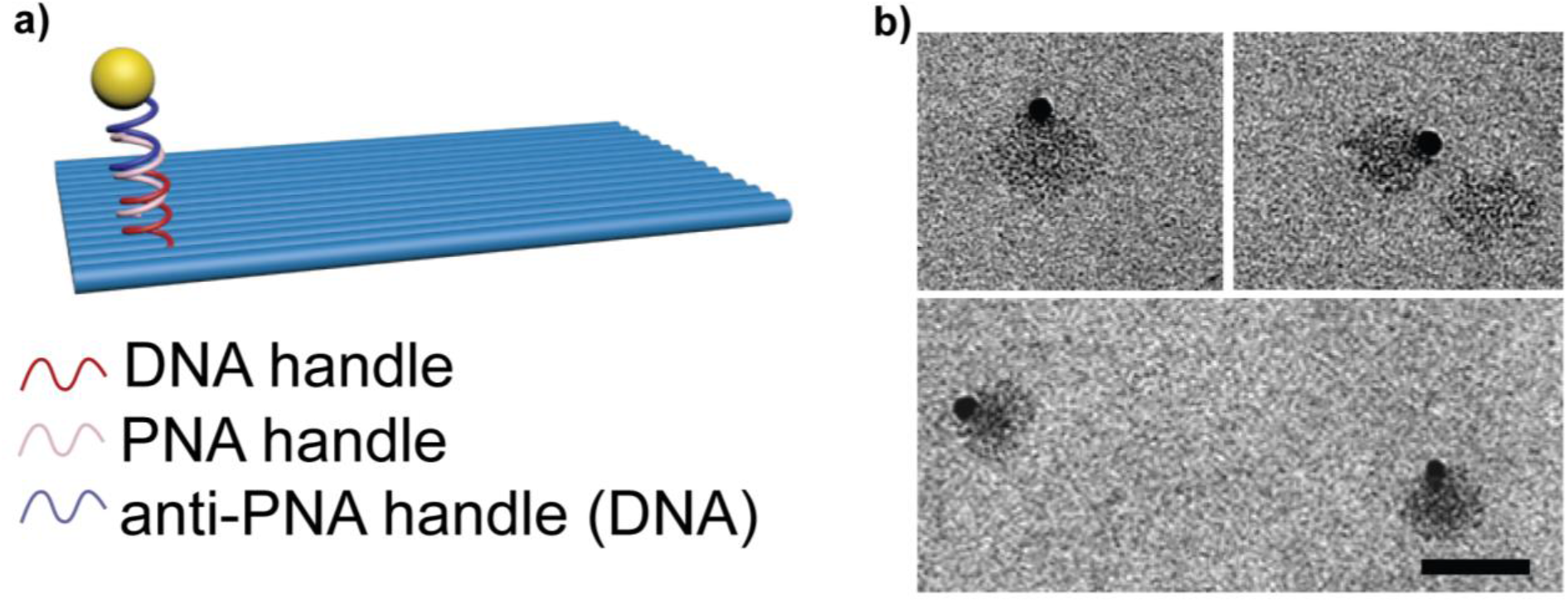
TEM images of silicified 1LS designed with protruding PNA handles, hybridized to 15 nm Au NPs. Scale bar is 100 nm. Structures were not stained.

**Table S1.**
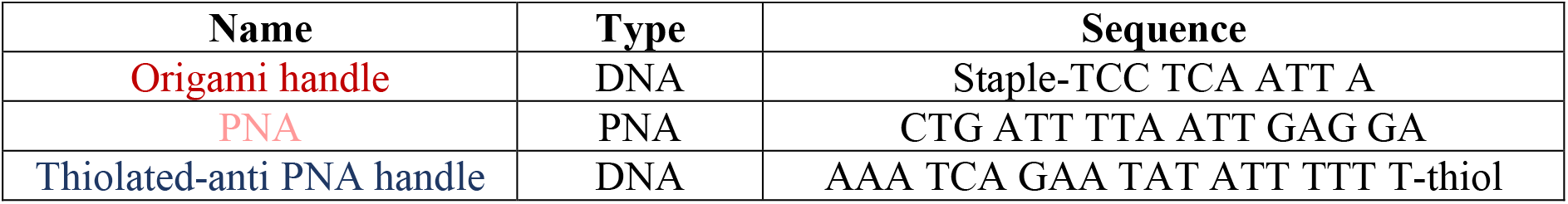
DNA and PNA Sequences

### Note S4: 4LB and silicified 4LB: TEM images

**Figure S7:**
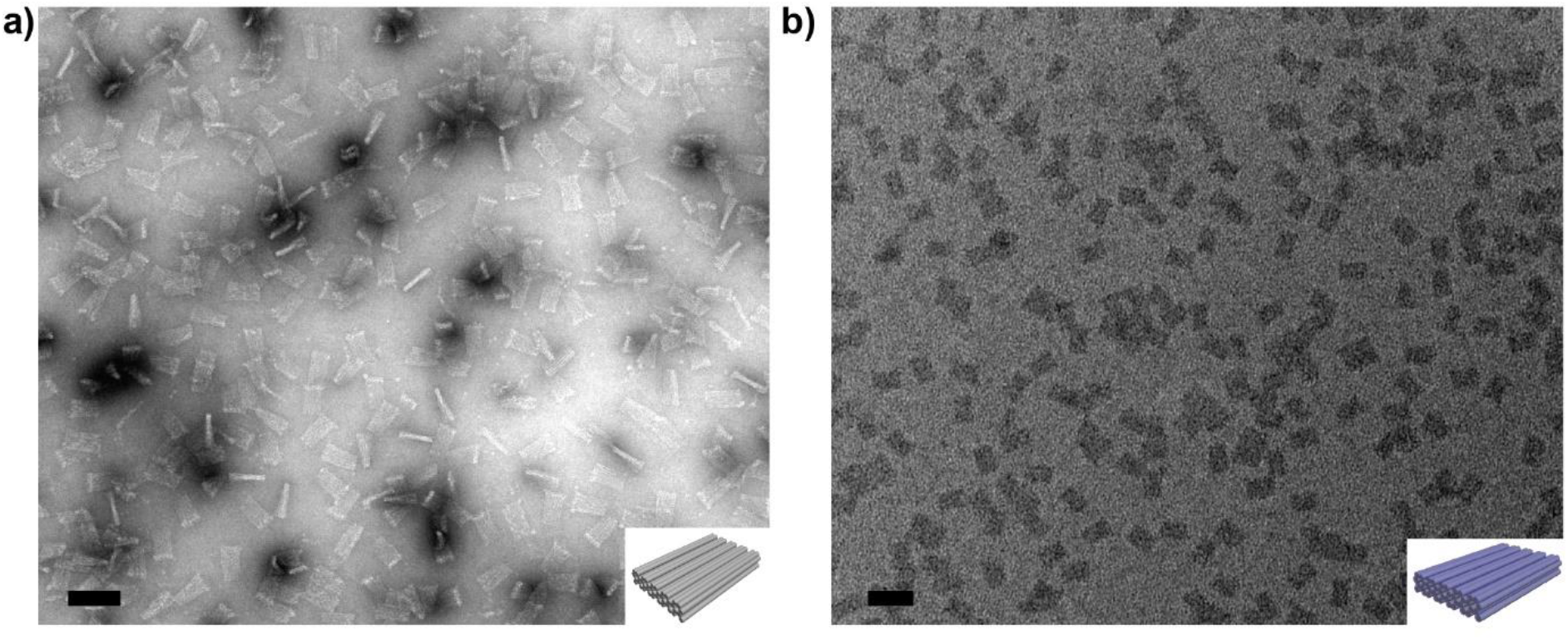
TEM micrographs of the 4LB a) before and b) after 4 h of silica growth in 3 mM MgCl_2_ at 21 °C using a revolving rotator at a DNA origami concentration of 200 nM. Bare structures were stained with uranyl formate, while silicified structures were not stained. Scale bars are 100 nm.

### Note S5: 24HB and silicified 24HB: TEM images

**Figure S8:**
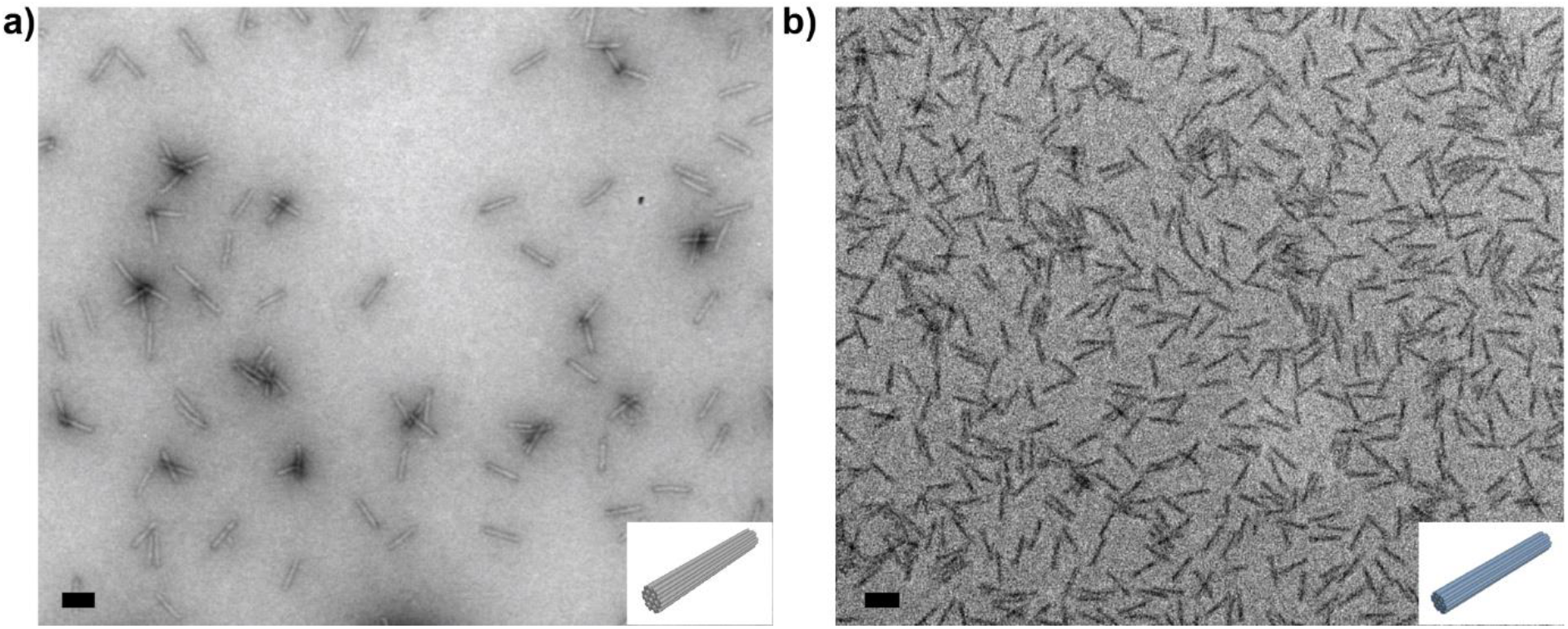
TEM micrographs of the 24HB before (a) and after (b) 4 h of silica growth in 3 mM MgCl_2_ at 21 °C using a revolving rotator at a DNA origami concentration of 200 nM. Bare structures were stained with uranyl formate, while silicified structures were not stained. Scale bars are 100 nm.

### Note S6: DNase stability (4LB & 24HB)

**Figure S9.**
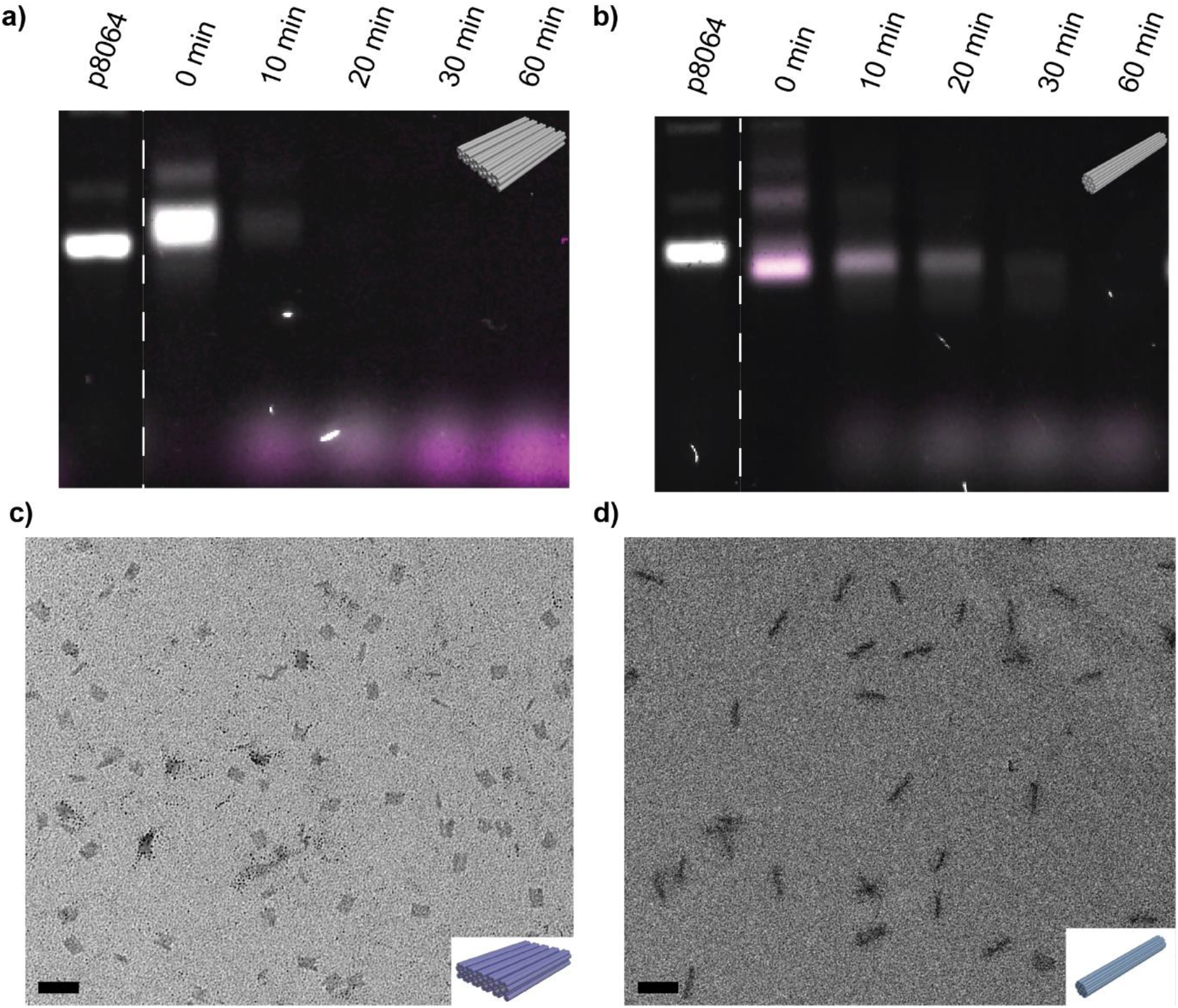
Agarose Gel electrophoresis of bare 4LB (a) and bare 24HB (b) after being incubated with DNase I for up to 60 min. The bare 4LB (a) already disintegrated after 10 min, while the bare 24HB was disintegrated after 20 mins. c) and d) show TEM micrographs of silicified 4LB (c) and silicified 24HB (d) after being incubated with DNase I for 3 h and 6 h respectively. Silica growth was done for 4 h in 3 mM MgCl2 at 21 °C using a revolving rotator at a concentration of 200 nM. Structures were not stained. Scale bars are 100 nm. Since the bare 4LB disintegrated faster than the 24HB and since our recent report found that 4LB structures did not silicify as well as 24HBs^[5]^, we incubated the silicified 4LB for a shorter amount of time with DNase I compared to the 24 HB.

### Note S7: Addressability of silicified 18HB

**Figure S10:**
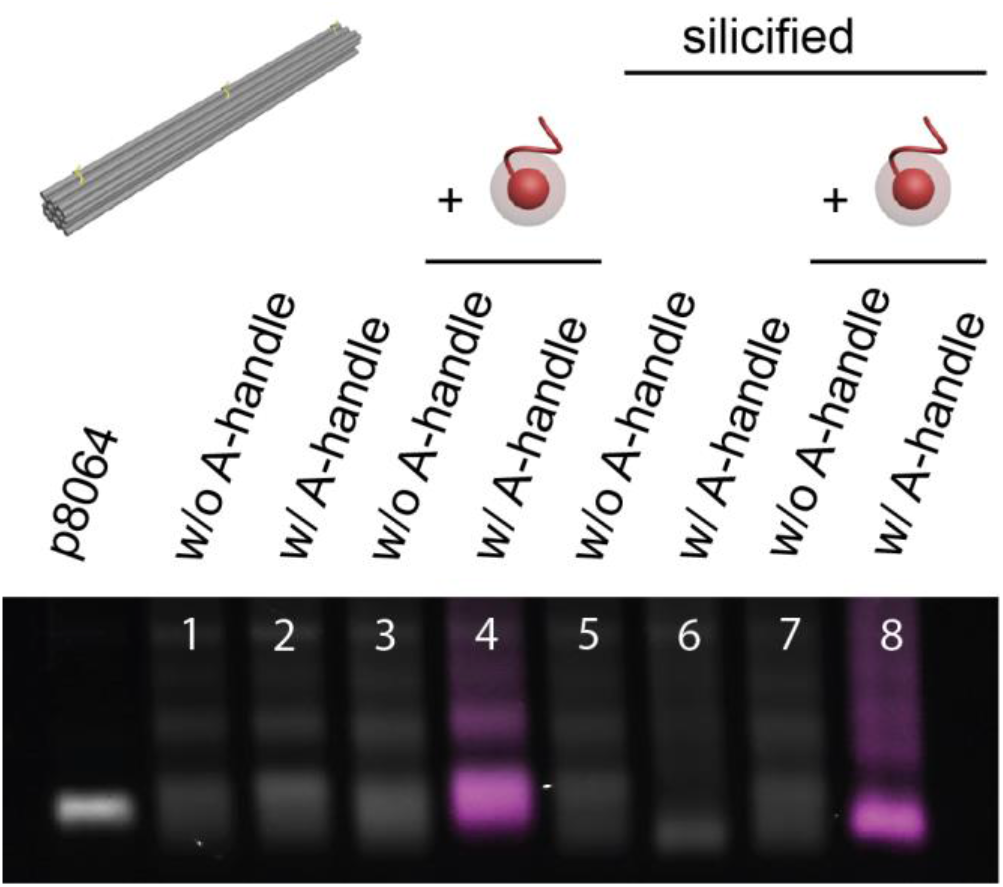
Agarose gel electrophoresis of the 18HB before (lanes 1 to 4) and after silicification (lanes 5 to 8) and with addition of the Cy5-anti-handle (lanes 3, 4, 7 and 8). Silicification was carried out for 4 h in 3 mM MgCl_2_ at 21 °C using a revolving rotator at a DNA origami concentration of 200 nM.

### Note S8: AFM imaging of surface immobilized SiO2 DNA origami

AFM images were leveled and background corrected prior to analysis. A homogenous silica shell growth is directly visible over the whole field of view in the corrected AFM images of bare (reference) and surface silicified 12HB nanostructures (**Figure S11a**, left and right, respectively). To further quantify the silica shell thickness, we extracted the pixel height distributions from the corrected AFM images. The height distributions show both a dominant peak around 0 nm height representing all background pixels and a second population shifted to higher z-values representing all pixels covered by DNA origami nanostructures (**Figure S11b**)). While the bare 12HB structures reveal a peak around 2.5 nm in height, the surface silicified 12HB structures showed a large shift to around 7.5 nm in height or a relative increase of around three-fold, resulting in a silica shell thickness of ca. 5 nm.

**Figure S11:**
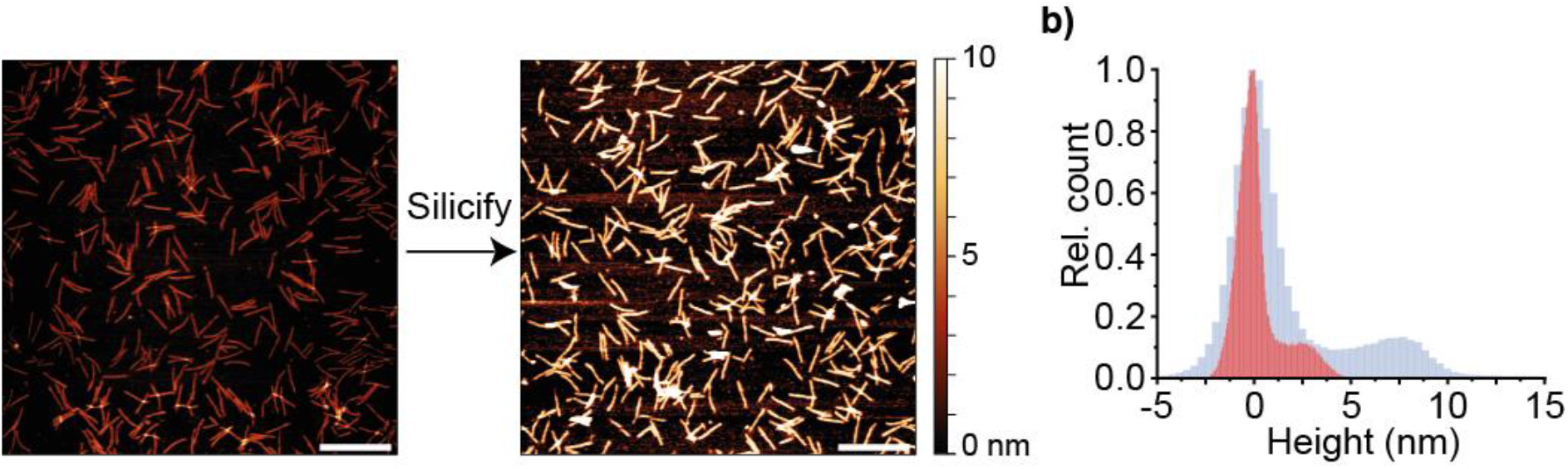
AFM images of the 12HB (a) before (left) and after silicification on the surface (right). Extracted height distributions of the 12HB (b) before (red) and after silicification (blue). Scale bars are 500 nm.

### Note S9: DNA PAINT imaging stability study of bare and silicified 12HB

To assess the stability of (silicified) 12HB nanostructures immobilized on the glass surface, they were incubated in degrading conditions (1×TAE buffer (without Mg^2+^) containing 1 mM EDTA) for 2 h.^[12]^ As expected, the bare 12HBs did not display the designed triple spot in DNA PAINT imaging anymore. Instead, mostly single spots were visible, indicating the structural collapse and degradation of the 12HB DNA origami (**Figure S12c**, left panel). Silicified 12HB on the contrary remained intact and still revealed the designed triple spot (**Figure S12c**, right panel), indicating significantly increased structural stability, due to the silica coating, even under harsh and otherwise degrading conditions. Picking individual labeling spots and extracting the binding kinetics leads to the dark time distributions given in **Figure S12d**. While the dark time distribution for the silicified 12HB after 2 h of incubation in degrading conditions did not change significantly (mean dark time of ca. 24.1 ± 9.8 s), the dark time distribution for the bare 12HB showed a significant shift to even shorter dark times (mean dark time of ca. 9.3 ± 3.1 s). While in the case of the silicified 12HB, the structure itself and the three labeling spots (each consisting of 6 DNA PAINT docking sites) stayed intact and thus the picking of individual spots leads to comparable results, the labeling situation in the case of the bare 12HB changed drastically during degradation: the initially three individual labeling spots with 90 nm distance collapsed into one single labeling spot consisting now of up to 18 individual DNA PAINT docking sites with unknown individual accessibilities. A shift to shorter dark times could thus be thus explained by the increase of DNA PAINT docking sites within one picked labeling spot.

**Figure S12:**
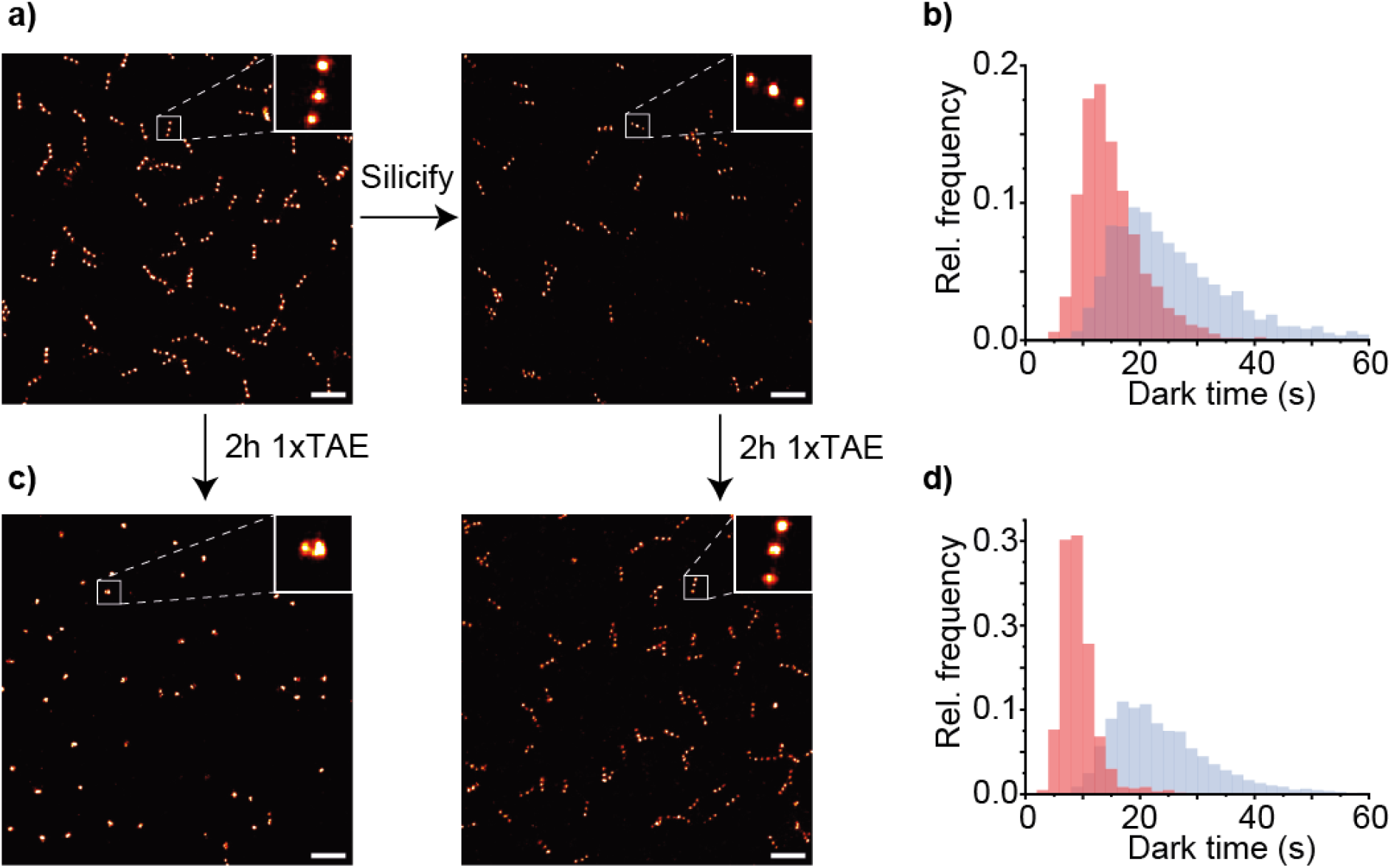
DNA PAINT images of the 12HB (a) before (left) and after silicification (right). (b) Extracted dark time histograms before (red) and after silicification (blue). DNA PAINT images (c) of the reference (left) and silicified (right) 12HB and corresponding extracted dark time histograms (d) of the bare (red) and the silicified 12HB (blue) after 2h incubation in 1×TAE buffer. Scale bars are 500 nm.

### Note S10: Dynamic DNA origami: bare and silicified 18HB TEM images and bending angle analysis

**Figure S13.**
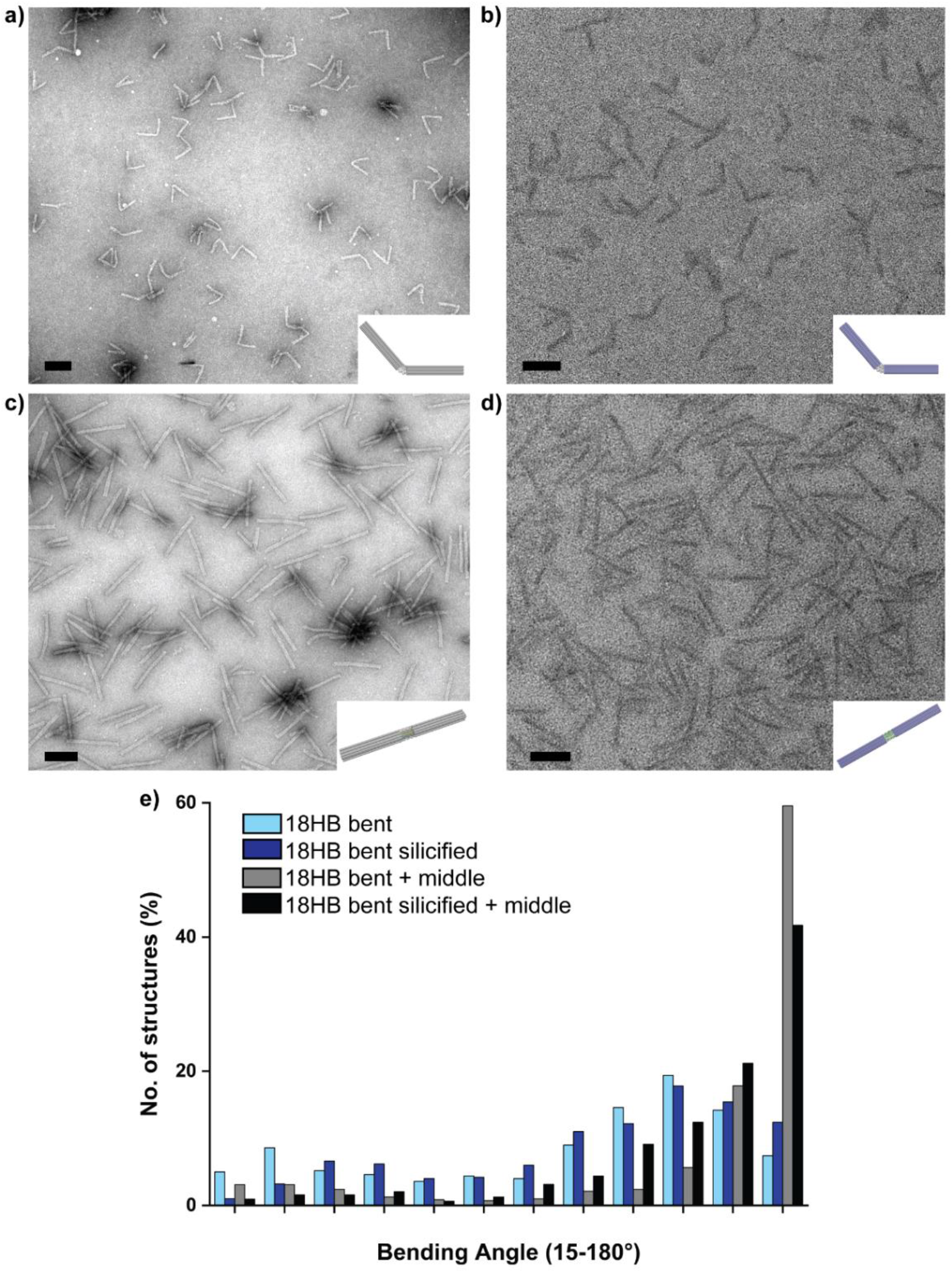
TEM micrographs of the bent 18HB before silicification (a), after silicification (b), after adding the middle staples (c) and after silicification and then adding the middle staples (d). Silicification was carried out for 4 h in 3 mM MgCl_2_ at 21 °C using a revolving rotator at a DNA origami concentration of 200 nM. Bare structures were stained with uranyl formate, while silicified structures were not stained. Scale bars are 100 nm. e) shows histograms of bending angle analysis of all four structures based on TEM images Analysis was carried out with ImageJ. More than 480 structures were analyzed for each condition. (Light blue, light grey: bare structures; dark blue, black: silicified structures).

### Note S11: Octahedral DNA origami (crystals)

**Figure S14.**
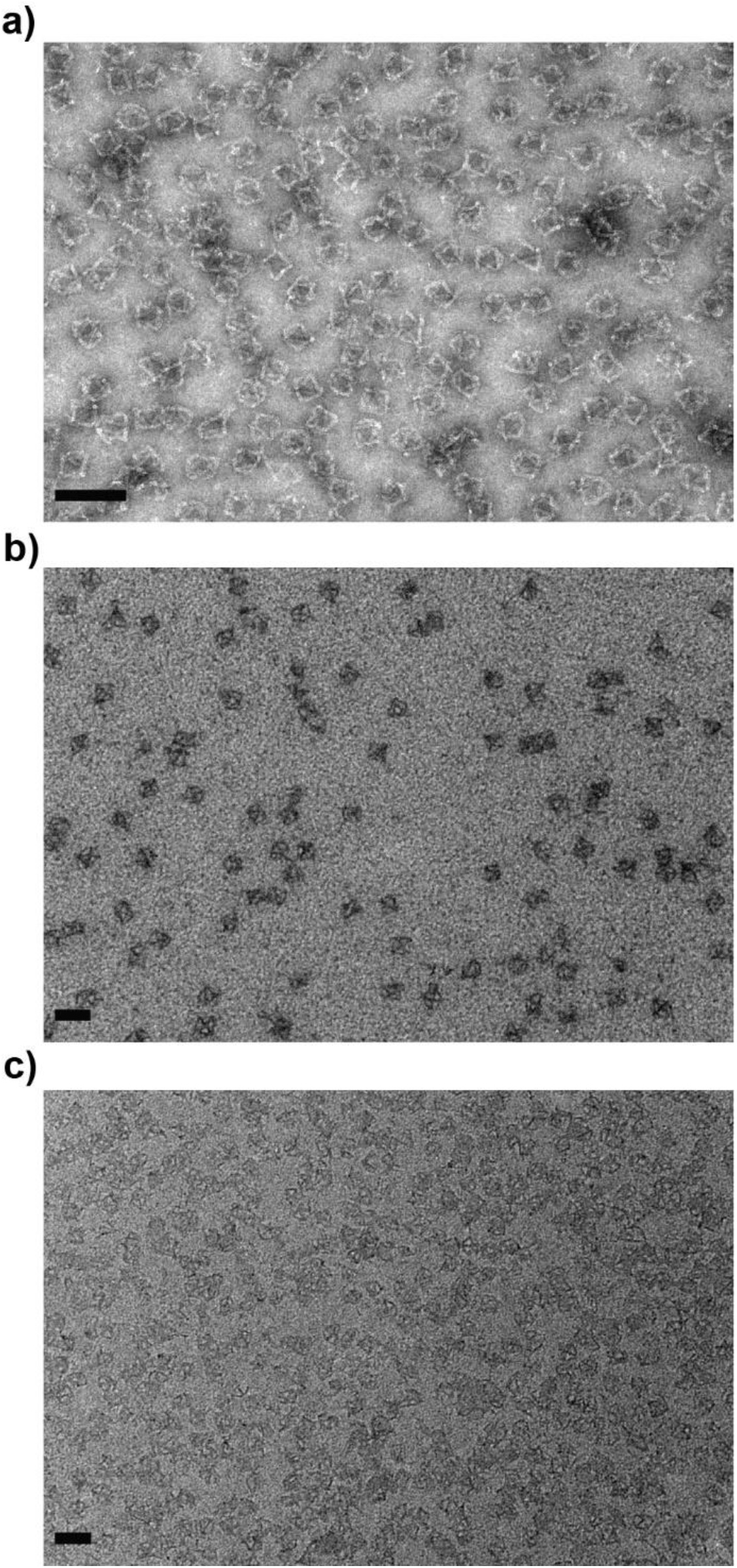
TEM images for bare (a) and silicified DNA origami octahedrons (b) and for silicified octahedrons after heating to 60 °C for 30 min (c). The images show a significantly increased rigidity and stability of the DNA origami nanostructures after silicification. Bare structures were stained with uranyl formate, while silicified structures were not stained. Scale bars are 100 nm.

**Figure S15.**
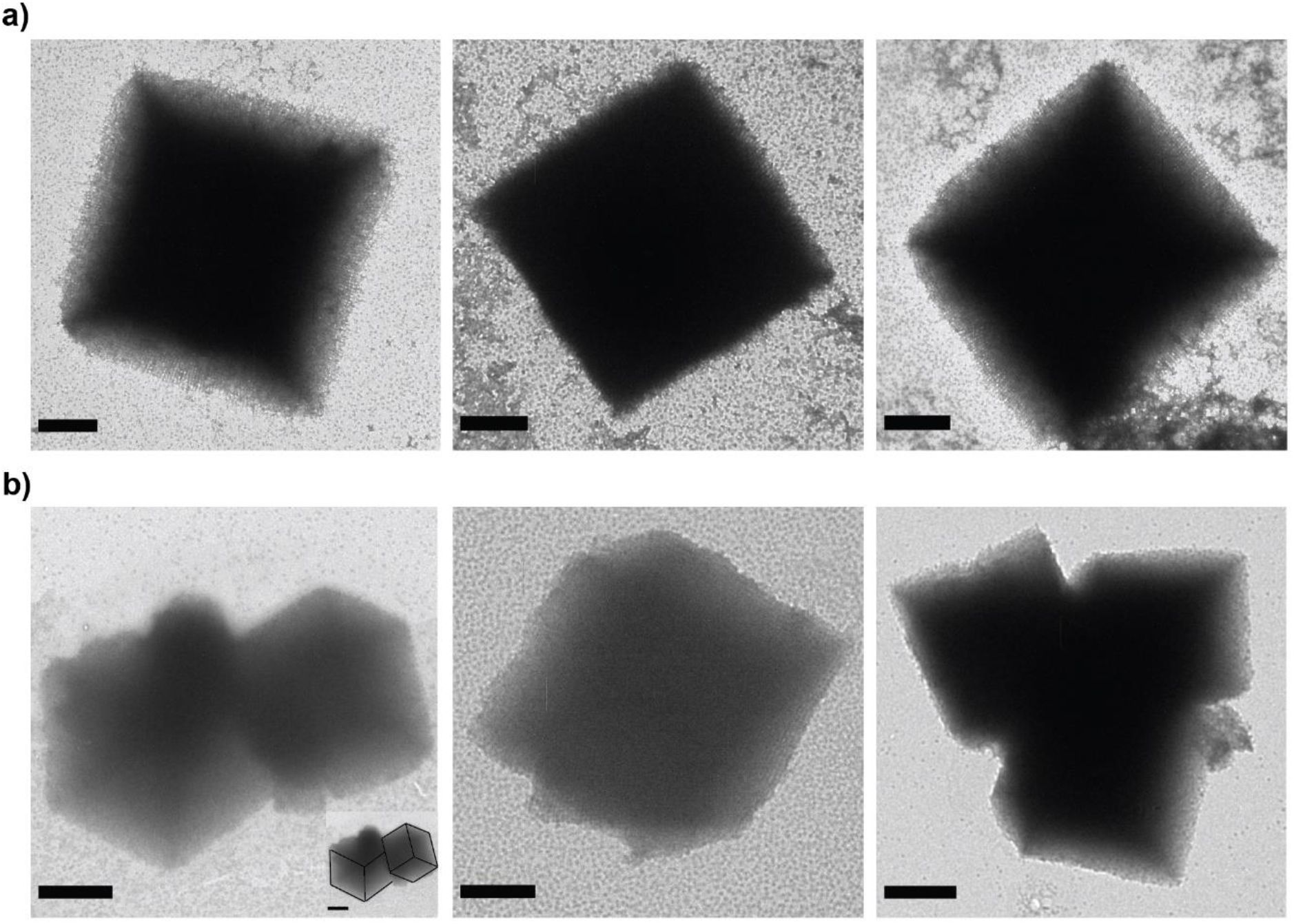
TEM images for cubic crystals made via polymerization of bare (a) and silicified DNA origami octahedrons (b). While the bare crystals almost always lie on one of the cubic faces, the crystal structures with silica can be deposited on their edges which makes their three-dimensional structure more visible and indicates an enhanced rigidity of the crystals. Bare structures were stained with uranyl formate, while silicified structures were not stained. Scale bars: 1 μm. Inset in b), left panel, shows the same image with a guide to the eye for 3D visualization.

### Note S12: Handle sequences

**Table S2:**
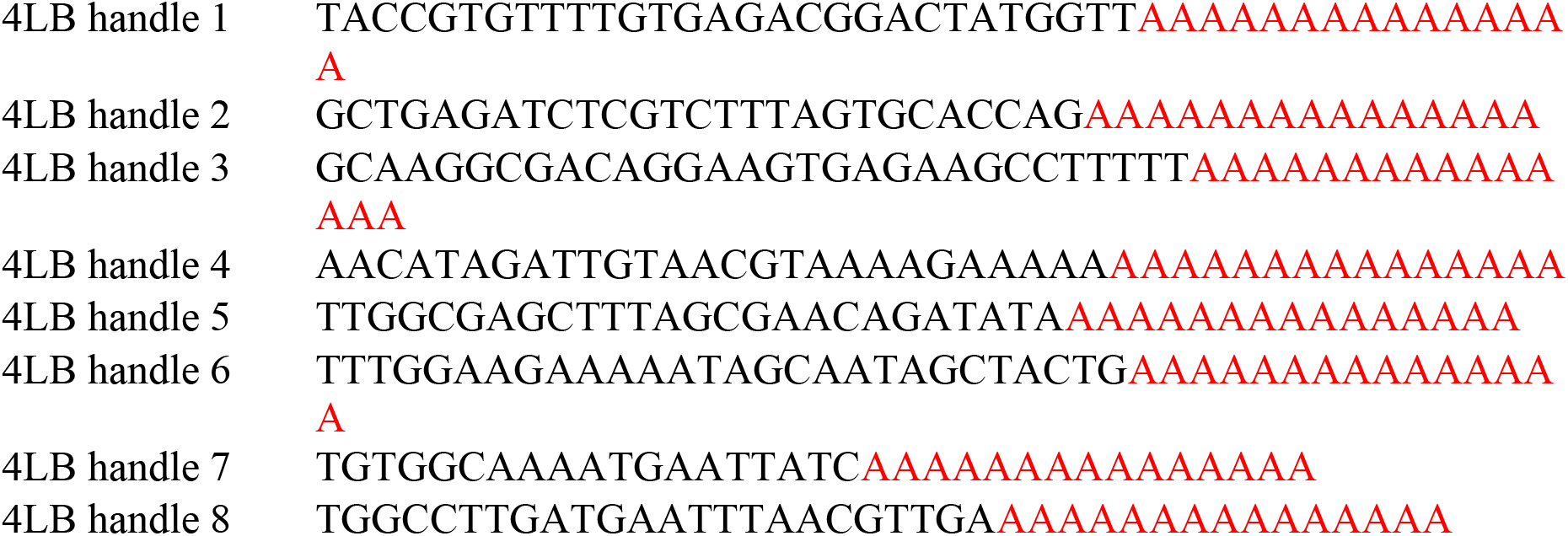
4LB handle sequences

**Table S3:**
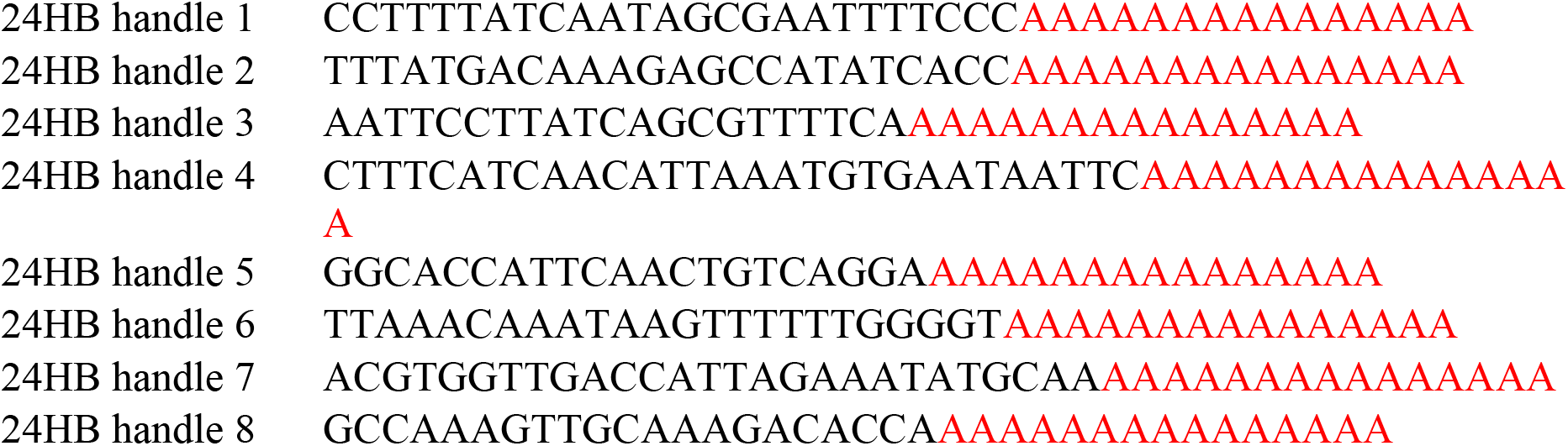
24HB handle sequences

**Table S4:**
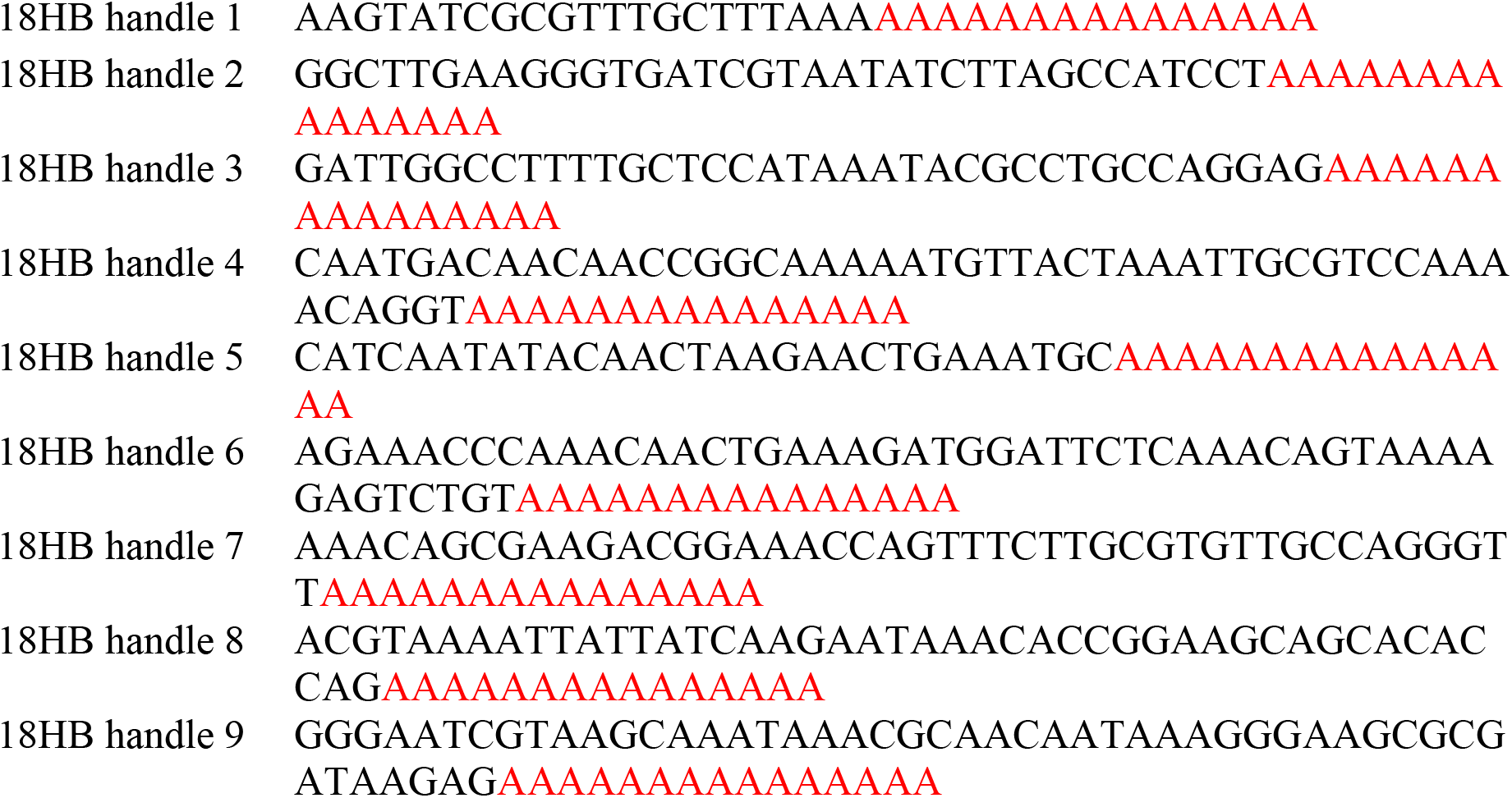
18HB handle sequences

**Table S5:**
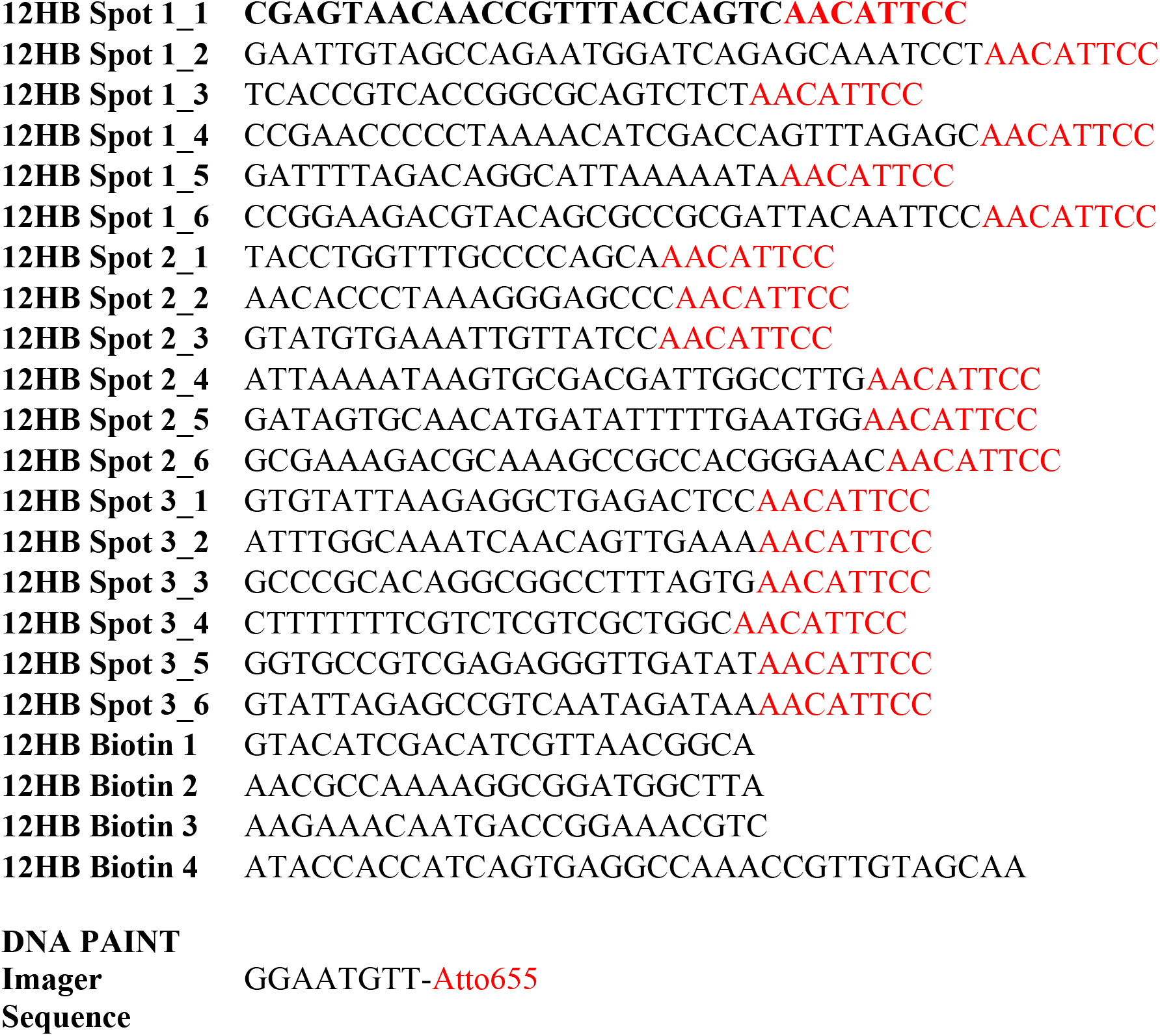
12HB handle sequences

**Table S6:**
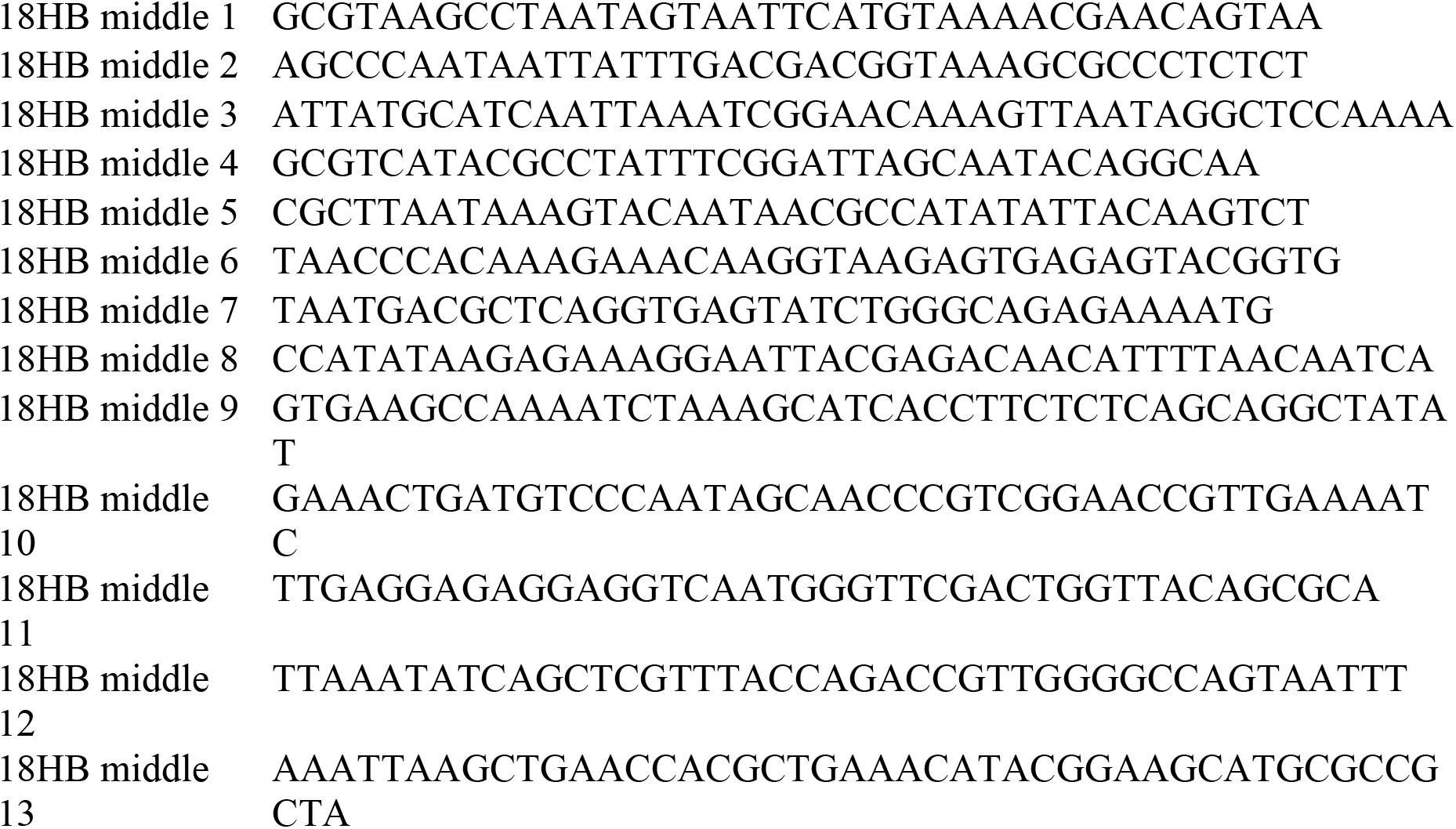
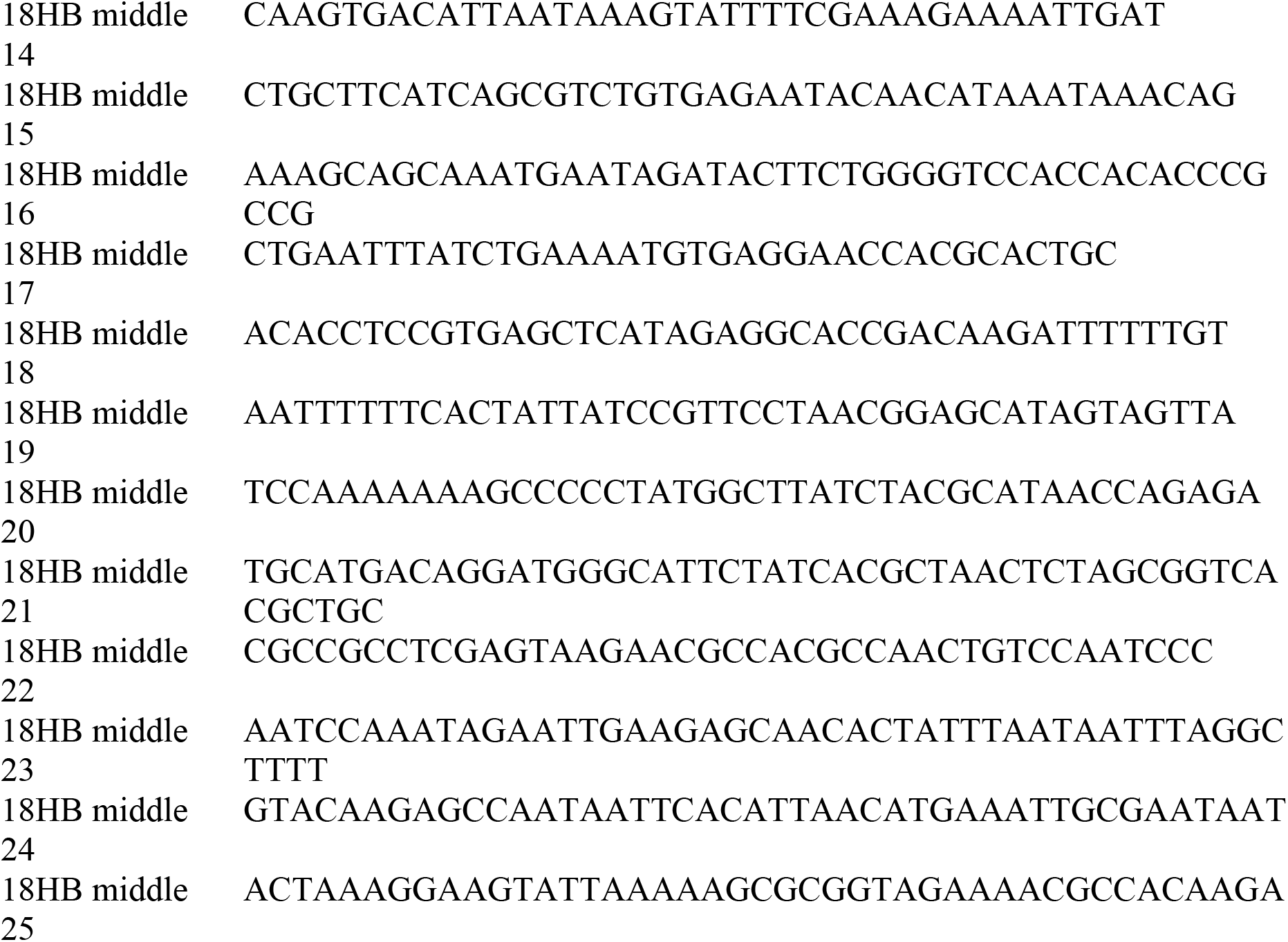
18HB middle bent sequences

